# *Staphylococcus aureus* secretes immunomodulatory RNA and DNA via membrane vesicles

**DOI:** 10.1101/2020.09.23.310318

**Authors:** Blanca V. Rodriguez, Meta J. Kuehn

**Affiliations:** Duke University, Department of Biochemistry, Durham, 27710, USA

## Abstract

Bacterial-derived RNA and DNA can function as ligands for intracellular receptor activation and induce downstream signaling to modulate the host response to bacterial infection. The mechanisms underlying the secretion of immunomodulatory RNA and DNA by pathogens such as *Staphylococcus aureus* and their delivery to intracellular host cell receptors are not well understood. Recently, extracellular membrane vesicle (MV) production has been proposed as a general secretion mechanism that could facilitate the delivery of functional bacterial nucleic acids into host cells. *S. aureus* produce membrane-bound, spherical, nano-sized, MVs packaged with a select array of bioactive macromolecules and they have been shown to play important roles in bacterial virulence and in immune modulation through the transmission of biologic signals to host cells. Here we show that *S. aureus* secretes RNA and DNA molecules that are mostly protected from degradation by their association with MVs. Importantly, we demonstrate that MVs can be delivered into cultured macrophage cells and subsequently stimulate a potent IFN-β response in recipient cells via activation of endosomal Toll-like receptors. These findings advance our understanding of the mechanisms by which bacterial nucleic acids traffic extracellularly to trigger the modulation of host immune responses.

## Introduction

Bacterial nucleic acids are well recognized as important pathogen associated molecular patterns (PAMPs) sensed by host innate immune cells during infection^1–3^. Nucleic acid-sensing receptors in eukaryotic cells have been characterized as complex intracellular sensors that must differentiate between ‘self’ and ‘intruding’ nucleic acids to mount a defensive host response against pathogens. Reports show that *Staphylococcus aureus* nucleic acids are important contributors to increased pro-inflammatory cytokine and Type I Interferon (IFN) signaling in host cells^4–7^. Studies have further identified that Toll-like receptors (TLRs) located within endosomal compartments of mammalian cells recognize the *S. aureus-derived* RNA and DNA molecules. A conserved sequence embedded within 23S ribosomal RNA has been identified as the minimal *S. aureus* RNA sequence required for TLR13 activation in mouse immune cells^7^. In human monocytes, TLR8 can sense *S. aureus* ssRNA degradation products containing a UR/URR RNA consensus motif, leading to IFN-ß production^8,9^. *S. aureus* tRNA was previously reported to induce Type I IFN production in peripheral blood mononuclear cells by activating TLR7^10^. These studies established a role for *S. aureus*-derived RNA in activating Type I IFN and cytokine production by transfection of purified *S. aureus* RNA into host cells or after treatment with heat-killed *S. aureus*. Additionally, a role for TLR9 in sensing staphylococcal DNA in murine bone marrow-derived dendritic cells was established^4^. Following treatment with lysates, or with viable and heat-killed bacteria, the authors showed a strong TLR9-mediated Type I IFN response to *S. aureus* DNA containing unmethylated CpG motifs^4^. Despite the growing body of literature showing the importance of *S. aureus* RNA and DNA in modulating host immune responses, there is a lack of mechanistic insight into how *S. aureus* secretes nucleic acids and how these molecules reach their cognate endosomal TLR receptors to alter host cell functions.

Recent studies in Gram-negative bacteria have revealed that extracellular vesicles produced by these organisms can potentially serve as delivery vehicles for RNA and DNA molecules to eukaryotic host cells^11–17^. Data from studies of *Pseudomonas aeruginosa* suggested that regulatory non-coding small RNAs (sRNAs) carried by these outer membrane vesicles (OMVs) may attenuate host innate immune responses in the mouse lung by repressing the expression of target host cell mRNAs that encode cytokines necessary for bacterial clearance^12^. In a separate study, uropathogenic *Escherichia coli* was found to deliver OMV-associated RNA into the cytoplasm and nucleus of human bladder epithelial cells *in vitro*^15^. Bitto et al also recently reported that chromosomal DNA fragments can be delivered to the nucleus of cultured lung epithelial cells by *P. aeruginosa* OMVs^17^. While such findings have catalyzed an interest in further understanding the functional roles of microbial vesicle-associated nucleic acids and how they modulate the immune response during infection^18–20^, these studies have not reported on the specific mechanisms by which OMV-associated RNA and DNA are taken up by host cells or provided evidence to show that OMV-associated nucleic acids interact directly with host cells to modulate the immune response. Additionally, studies of the extracellular vesicles produced by Gram-positive bacteria (termed membrane vesicles, MVs) have primarily focused on their proteinaceous cargo, with limited characterization of the MV-associated RNA and DNA content^21–24^. Therefore, although MVs are increasingly recognized as key mediators in the dynamic crosstalk between the pathogen and the host^11,25–28^, the functional relevance and transfer of MV-associated RNA and DNA cargo from Gram-positive bacteria to host immune cells remains to be defined.

MVs are a heterogeneous population of spherical, bioactive nanoparticles that range from 20 to 300 nm in diameter, secreted naturally by many Gram-positive bacteria over the course of their lifecycle^25^. Originating from the cytoplasmic membrane, MVs contain an array of cellular components that include virulence factors, phospholipids, polysaccharides, small molecules, and lipoproteins. Biochemical and proteomic analyses have shown that the molecular cargo of MVs includes RNA, DNA, and cytoplasmic proteins that lack export signals^21,22,25,29–33^. MV cargo can be associated to the surface of vesicles or encapsulated within the MV lumen^27^. Importantly, MVs were shown to stabilize and protect their protein cargo from extracellular proteolytic digestion and likely also protect their nucleic acid cargo from degradation by nucleases present in the extracellular space. This feature enables MVs to deliver a selection of protected, biologically active macromolecules to host cells both proximal and distal from the site of bacterial infection^27^. Generally, MVs are internalized by host cells via endocytosis, macropinocytosis, or phagocytosis—which facilitates the delivery of bacterial-derived immunomodulatory cargo to mammalian host cells in a coordinated fashion^27,34,35^. Such reports position MVs as a likely, non-canonical secretion mechanism for bacterial nucleic acids to gain access to the interior of host cells and to play a functional role in the host-pathogen interface.

DNA and RNA secretion has been characterized for some important pathogens. For instance, the release of Gram-negative extracellular DNA by the type IV secretion system has been well-described^36^. This mechanism is largely dependent on direct cell-to-cell contact, although contact-independent export has been observed in *H. pylori* and *N. gonorrhoeae.* In contrast to DNA release, RNA secretion mechanisms remain largely unexplored. Recent studies of the intracellular pathogens *Mycobacterium tuberculosis* and *Listeria monocytogenes* have suggested the existence of MV-independent RNA secretion mechanisms that elicit a potent IFN-ß response in host cells^37–39^. Cheng and Shorey showed that *M. tuberculosis* releases RNA into the cytosol of infected macrophages in a SecA2 and EsX-1 secretion system-dependent manner, however they only speculated that potential RNA-binding proteins could also shuttle bound RNA molecules via the SecA2 system into the extracellular milieu^37^. Frantz and colleagues recently showed that *L. monocytogenes* secretes both “naked” and MV-contained sRNAs that are potent IFN-ß inducers in macrophages and in HEK293 cells^38^. This group focused on characterizing specific “naked” sRNAs present in the non-MV containing supernatant fraction that also induced high levels of IFN-ß but did not examine the secretion mechanism for the RNA. Notably, no specialized nucleic acid secretion systems have been reported for *S. aureus.*

The present study aims to elucidate the nature of the association between nucleic acids and MVs produced by the human pathogen *S. aureus.* We also seek to analyze the immunostimulatory potential of MV-associated RNA and DNA, and to evaluate receptor-mediated recognition of MV-associated RNA and DNA molecules by innate immune cells. Consistent with the hypothesis that nuclease protected MV-associated RNA and DNA molecules induce Type I IFN production in mouse immune cells, we found that endosomal *TLR3*^-/-^, *TLR7*^-/-^, and *TLR9*^-/-^ macrophages expressed significantly reduced IFN-β mRNA in response to nuclease-treated MVs relative to their wild-type counterparts. We demonstrated that uptake of MVs followed by endosomal acidification and maturation are critical steps that lead to a potent IFN-β response in macrophages. Collectively, these findings point to a mechanism whereby MVs facilitate the transfer of bacterial RNA and DNA molecules into recipient cells.

## Results

### *S. aureus* Newman strain secretes MVs containing nucleic acids during normal growth conditions

MVs in cell-free *S. aureus* Newman culture supernatants were separated from protein aggregates, cell debris, and non-membranous soluble proteins using Optiprep gradient density centrifugation as illustrated in Fig. S1. A total of 6 fractions were collected and the protein, lipid, and nucleic acid content of each fraction was measured. The lowest density fractions (F1, F2, and F3) contained the highest protein and lipid content as determined by Bradford and FM4-64 assays, suggesting the presence of the MVs (Fig. 1A). Notably, these fractions also contained the highest DNA and RNA content as determined using fluorescence-based dyes (Fig 1A). The presence of MVs in the fractions was assessed using negative-stain TEM analysis (Fig. 1B and S2). In accord with our proteolipid quantification analysis, the lower density fractions F1-F3 were highly enriched with MVs, whereas few vesicles were identified in the higher density fractions (Fig. 1C). Morphologically, MVs of average smaller diameter were distributed to the lower density fractions (F1-F3), in contrast to higher density fractions (F4-F6) where larger sized MVs were observed. Consistent with previous data regarding *S. aureus* MVs^25,33,40–44^, our size distribution analysis of the pooled MV fractions F1-F3 showed a size distribution range from ~20 nm to ~300 nm in diameter (Fig. 1D), with an average diameter of 68.5 nm. Due to the abundance of MVs in the lower density fractions, F1-F3 were pooled and used for the MV characterization and functional experiments described hereafter.

**Figure 1.**
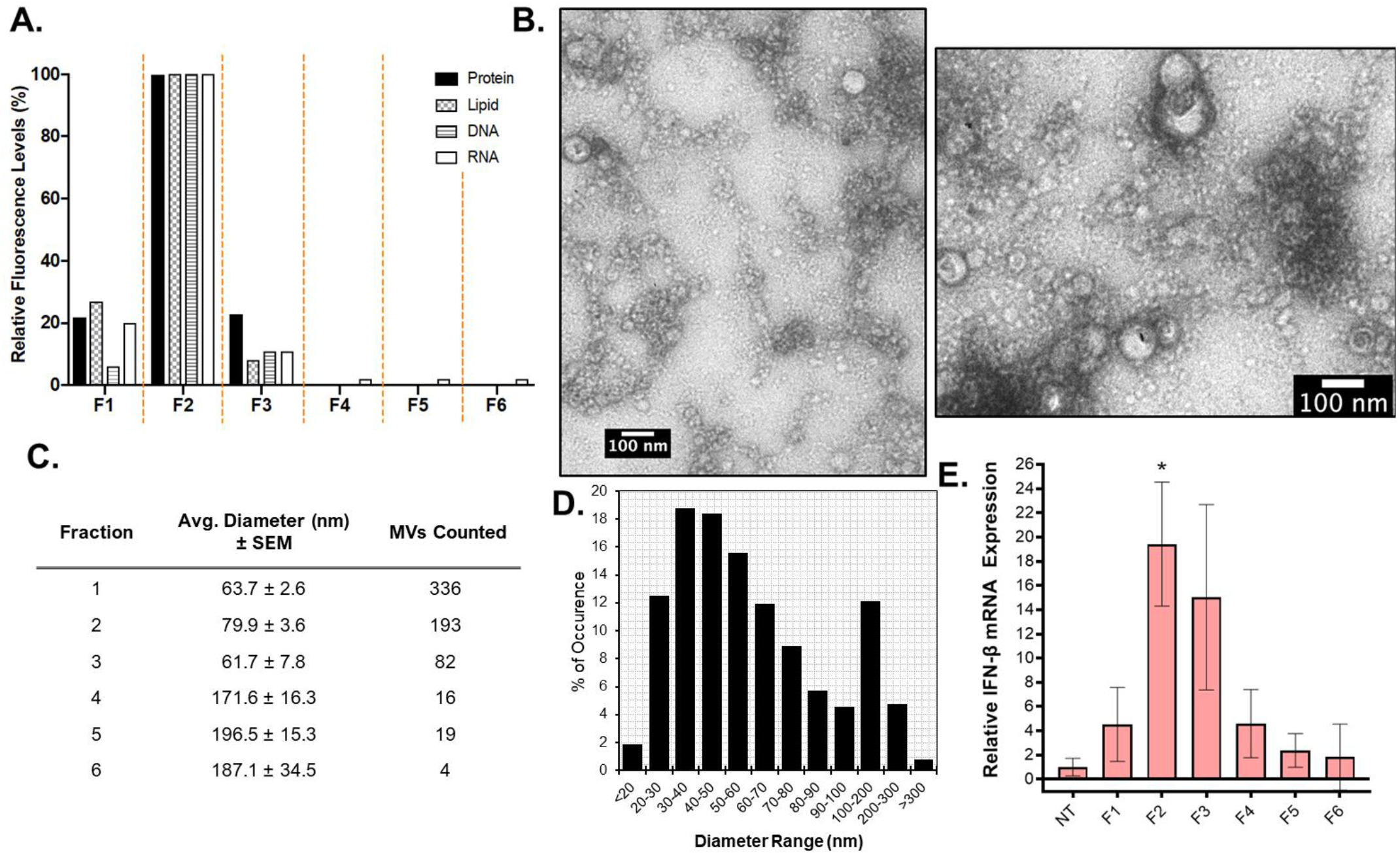
*S. aureus* releases MVs enriched in proteins and nucleic acids. (A) All six Optiprep fractions (F1-F6) were analyzed for protein, lipid, DNA, and RNA content using Bradford, FM4-64, SYTOX Green, and SYTO RNASelect, respectively. Relative fluorescence values shown were normalized to 100% in the peak fraction for each method. (B) Negative-stain TEM was used to image purified MVs. Representative images of fraction 2 are shown. (C) The size and number of MVs imaged in each fraction from a 10 μL sample in a negative-stained copper grid was determined using Fiji. The average diameter ± standard error of the mean (SEM) of MVs counted in each fraction are tabulated. (D) Size distribution analysis of pooled MV fractions F1-F3 is presented as MV frequency of occurrence by size range. (E) q-PCR was used to determine the IFN-ß gene expression levels in RAW264.7 cells after exposure to 0.5 μg/mL of each fraction for 3 h. IFN-ß mRNA expression was normalized to ß-actin mRNA expression. Results are representative of three independent experiments. Statistical analysis was performed by one-way ANOVA with Tukey’s multiple comparison test (n.s = not significant; *p ≤ 0.05).

MV production could potentially serve as an active secretion system for nucleic acids produced by viable, intact *S. aureus* cells; however, it is conceivable that nucleic acids could be released through spontaneous phage-dependent or independent cell lysis^29–31,45,46^ and that such nucleic acids could become associated with MVs after MVs are released into the extracellular space. The current state of knowledge regarding bacterial MV formation and the possible routes by which cytoplasmic material, such as DNA, could be loaded into MVs has recently been summarized by Toyofuku and colleagues^47^. So far, two independent groups have provided evidence for the formation of certain MV subpopulations through routes that involve weakening of the peptidoglycan layer by cell wall-degrading enzymatic activity^29,30^. In *Bacillus subtilis*, prophage-encoded endolysins were activated by genotoxic stress leading to the release of a sub-population of MVs from dying *B. subtilis* cells^29^. In *S. aureus* JE2, production of certain MV subpopulations was promoted by the coordinated activity of phenol-soluble modulins and peptidoglycan-targeting autolysins^30^. Additional routes of MV formation in lysogenic and non-lysogenic *S. aureus* strains were shown to be induced by a variety of different antibiotics, which had consequences on the DNA cargo packaged in MVs^31^. Taking into account all of these findings, we deliberately aimed to reduce the amount of exogenous DNA released by broken cells by purifying MVs harvested from *S. aureus* cultures grown to mid-log phase (OD_600_ of ~1.0, Fig. S3A). We then verified that at this growth stage there were minimal levels of cell lysis by using a SYTOX Green assay and a LIVE/DEAD BacLight Bacterial Viability Kit to probe membrane integrity and cell viability (Fig. S3 B-D). Based on these results, MVs purified for this study were collected from mid-exponential phase cultures. It is important to note however, that spontaneous cell lysis, even at minimal levels could potentially still contribute to the association of DNA in MV preparations.

While cell death was shown to be minimal, we also considered the possibility that MVs could co-purify with phages released by *S. aureus* Newman, a known lysogenic strain^48^. If MVs samples contain co-purified phages, this could confound our analysis of *bona fide* MV-associated DNA. We previously showed that distinct T4 phages can be imaged by TEM in association with *E. coli* OMVs after co-incubation, suggesting that both MV morphology and the structure of exogenous phages remain relatively stable after co-incubation^49^. Additionally, antibiotic-induced MVs purified from a lysogenic strain of *S. aureus* were shown to co-purify with filamentous phage-like structures as imaged by TEM^31^. To our knowledge however, there is no evidence that complete and intact phages can be recovered from non-induced *S. aureus* MV samples after purification by Optiprep gradient centrifugation. Indeed, after density gradient centrifugation no filamentous or tail-like structures were evident in the *S. aureus* Newman MV TEM micrographs presented herein (Fig. 1B and S2). While we cannot rule out the possibility that phages may co-purify with MVs, even if they are present after Optiprep purification, they may be present in extremely low abundance.

We also considered the possibility that, despite harvesting MVs from only mid-log cultures, RNA derived from cell lysis may co-purify with MVs or adhere to the surface of MVs during the MV purification process and confound interpretation of our data. To address this concern, we first tested whether RNA that we added exogenously to MVs could be removed from the MVs with RNase treatment. The exogenously added *S. aureus* cellular RNA remained stable in MV samples after 30 min of incubation, however, RNase A treatment of the exogenous RNA-MV mixtures completely digested the exogenously added RNA from the samples (Fig. S4A). This indicated that in our future experiments using purified MVs, RNase A treatment would efficiently remove any exogenous RNA that had become associated with the MVs during culture growth or MV purification. Similar results were observed when purified MVs were incubated with exogenously added DNA in the form of the standard DNA ladder (Fig. S4B). Next, we characterized how readily exogenous RNA adsorbed to MVs by incubating cellular RNA with purified MVs for 30 min, subjecting the MV-RNA mixtures to ultracentrifugation, washing unbound molecules from MVs, and monitoring the binding of the exogenous RNA to the MVs using agarose gel electrophoresis. In this case, we did not detect the intact exogenously added RNA in association with MVs after the addition of a wash step (Fig. S4C). In addition, exogenous RNA recovered from the MV-free supernatant consisted predominantly of small RNA, indicative of RNA degradation. This could mean that the MV purification process of high-speed ultracentrifugation can lead to degradation of unprotected RNA molecules. Based on the fact that exogenously added RNA did not appear to adsorb to purified MVs, we propose that the native MV-associated RNA cargo is likely packaged into MVs during MV formation. While there is a possibility that RNA degradation fragments from the exogenously added RNA could become associated with MVs during ultracentrifugation through specific or non-specific binding, no additional bands that were not already present in the MV-only samples were detected (Fig. S4C).

A final consideration was the possibility that some of the RNA that is detected in association with MVs could be carried over from the rich culture media in which *S. aureus* is grown. In fact, Ghosal *et al*. previously identified RNA molecules of yeast origin in *E. coli* OMVs, likely originating from the Luria-Bertani media used to culture the bacteria^50^. Consequently, for the MV preparations used throughout this study, we treated the growth media with RNase A prior to sterilization and inoculation, a process that did not drastically affect the growth kinetics of *S. aureus* (Fig. S3A). Altogether, these data suggest that nucleic acid bearing-MVs harvested from *S. aureus* cultures grown to mid-log phase are mostly secreted by actively growing cells.

### Cultured murine macrophages produce significant Interferon-ß mRNA in response to MV stimulation

After establishing that *S. aureus* Newman strain secretes nucleic acids in association with MVs, we set out to investigate whether purified MVs induce Type I Interferon mRNA expression in innate immune cells as has been previously reported for *S. aureus* derived nucleic acids^4–7^. Both time-dependent and dose-dependent responses were assessed in RAW264.7 macrophage cells. Time-course experiments showed significant IFN-β mRNA induction within 1 hour of stimulation with MVs (5 μg/mL, by protein) and a peak expression of IFN-β mRNA at 3 hours (Fig. 2A). These results were in agreement with previous studies that reported peak IFN-β mRNA expression between 2-4 hours by airway epithelial cells and bone marrow-derived macrophages post-infection with *S. aureus* bacteria^5,51,52^. We also observed a significant expression of IFN-β relative to untreated macrophages with as low as 1 μg/mL MVs and a significantly increased, dose-dependent IFN-β response with 5 μg/mL MVs (Fig. 2B). Based on these results, MV-stimulated macrophages were harvested for RNA extraction 3 hours after exposure to MVs (5 μg/mL), unless indicated otherwise. We also found that macrophages preferentially induced IFN-β mRNA expression in response to MV stimulation, whereas MVs did not induce IFN-***α*** expression (Fig. 2C). This result is consistent with previous reports that showed a marked capacity for macrophages to produce IFN-β in response to *S. aureus* bacterial infection, whereas IFN-α is generally secreted by plasmacytoid dendritic cells after bacterial infection^5,53^.

**Figure 2.**
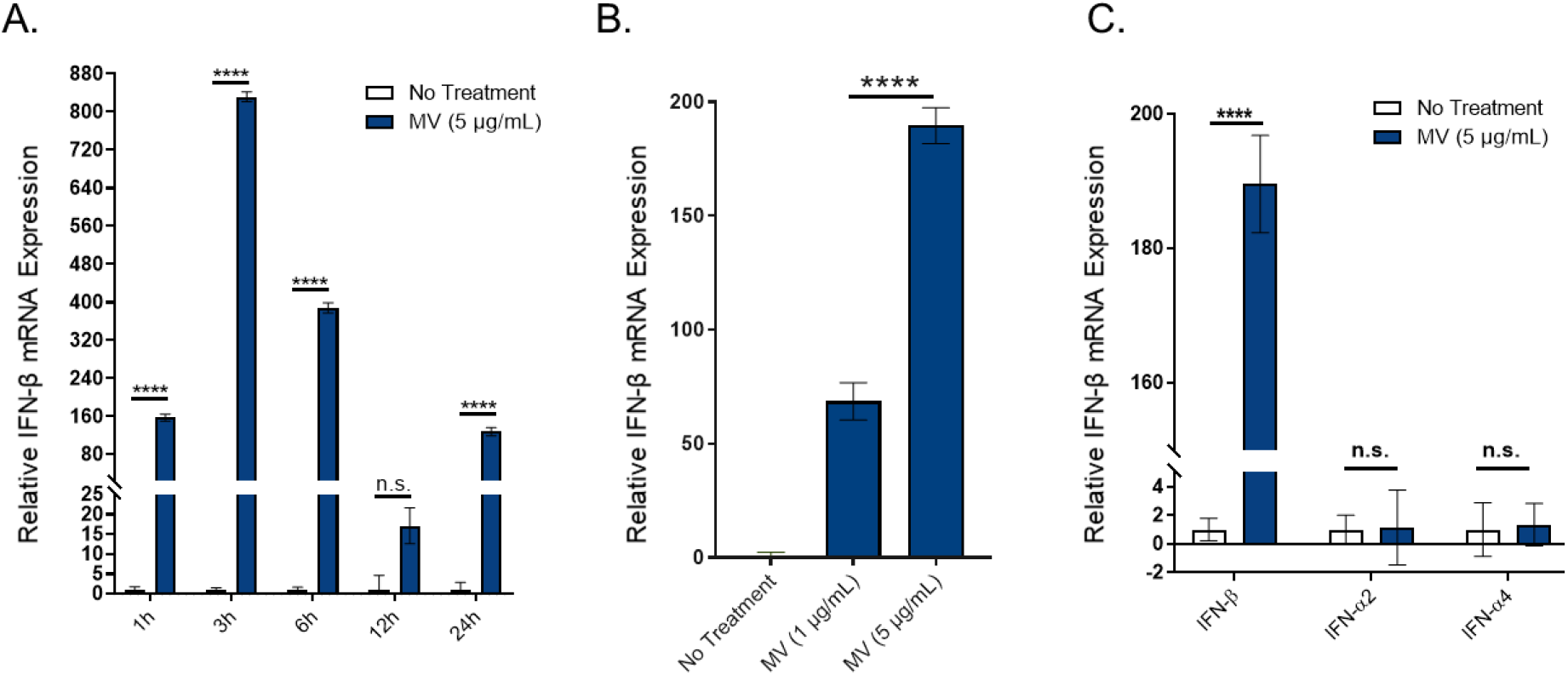
Purified *S. aureus* MVs induce significant IFN-β mRNA expression in cultured murine macrophages. (A) Time-course analysis of RAW264.7 cells treated with MVs (5 μg/mL total protein) over a 24 h time period. Macrophages were homogenized at the indicated times points for RNA extraction and analyzed for β-actin and IFN-β mRNA expression using qPCR. The relative fold induction of a sample was calculated relative to the no treatment control sample using the Ct method. β-actin was used to normalize IFN-β mRNA expression in all samples. (B) Macrophages were stimulated with the indicated doses of *S. aureus* MVs for 3 h and the IFN-β mRNA expression relative to β-actin was determined by qPCR. (C) Macrophages were incubated with no treatment controls or with MVs (5 μg/mL) and IFN-α2, IFN-α4, or IFN-β mRNA expression was measured using qPCR. The data is represented as mean SEM, n = 3. Profiles are representative of at least two independent experiments. Statistical analysis was performed by one-way ANOVA with Tukey’s multiple comparison test (****p < 0.0001).

Because these experiments were performed using pooled MV fractions, we wondered whether the highest MV activity—in terms of IFN-β mRNA induction—corresponded to the 3 low-density fractions enriched with MVs and nucleic acids. As shown in Figure 1E, when we stimulated macrophages with equal total protein concentration (0.5 μg/mL) of each fraction, we observed significant IFN-β expression in response to F2, while no significant IFN-β mRNA was detected in response to the remaining fractions. These results show that lower density fractions do in fact contain immunomodulatory MVs and that normalizing the MV dosage by total protein concentration—as is standard in the extracellular vesicle field—is a convenient metric to use to normalize the amount of MVs that are added to macrophages for stimulation assays between experiments. Collectively, these data show that *S. aureus* MVs induce strong IFN-β mRNA expression in cultured macrophage cells in a dose- and time-dependent manner.

While *S. aureus* does not produce lipopolysaccharide (LPS), endotoxins are ubiquitous environmental contaminants commonly found in the laboratory setting^54^. LPS acts as a strong inflammatory mediator in mammalian cells through activation of the surface-exposed receptor TLR4^55^. LPS-activated cells mount an immune response consisting of pro-inflammatory cytokine release and Type I Interferon production. In order to address the possibility that the IFN-ß mRNA macrophage response to MVs was due to endotoxin contamination, we evaluated the presence of LPS in *S. aureus* MV samples by SDS-PAGE and Western blot analysis. LPS was not detected in any of the MV samples used for this study as noted by the absence of the distinctive LPS laddered banding pattern in MVs after staining of the SDS-PAGE gel using the Pro-Q Emerald LPS Gel Stain (Fig. S5A). The high molecular weight bands detected by the LPS stain in MV samples were also observed using the SYPRO Ruby Protein Gel Stain, indicating that those bands are glycoproteins and not LPS (Fig. S5B). Additionally, we show that LPS was not detected in *S. aureus* MVs by Western blot analysis using goat polyclonal antibodies directed against the endotoxin region of LPS. Non-specific binding of anti-LPS antibodies to immunoglobulin-binding proteins in *S. aureus* MVs (such as SpA and/or Sbi) was responsible for the bands detected in MV samples (Fig. S5C). This was confirmed by complete removal of non-specific bands in MVs after proteinase K treatment, whereas bands corresponding to the LPS standard were unaffected by proteinase K (Fig. S5D).

Staphylococcal protein A (SpA) was previously shown to induce IFN-ß mRNA expression in human airway epithelial cells following endocytosis of the purified protein^52^. Furthermore, proteomic analyses have revealed that *S. aureus* MVs contain SpA^30,40,41^ and our own work suggests that IgG binding proteins, such as SpA are associated with MVs (Fig. S5D). Therefore, we examined whether SpA could contribute to the IFN-ß mRNA response of macrophages to MVs. We first established that Newman Δ*spa* produces MVs at late log-phase as confirmed by TEM imaging of MVs collected at OD_600_ of 0.9 (Fig. S6A & B). The protein profile of Δ*spa* MVs was analyzed by SDS-PAGE (Fig. S6C). We show that wild-type and Δ*spa* MVs contain some similar protein bands, while other proteins appear to only be packaged into either wild-type or Δ*spa* MVs (Fig. S6C). By qPCR, we show that MVs produced by *S. aureus Δspa* retain their ability to induce significant IFN-ß mRNA in cultured macrophages, suggesting that other macromolecules, such as nucleic acids could also be involved in eliciting the IFN-ß response (Fig. S6D). However*, Δspa* MVs induce IFN-ß mRNA in macrophages at significantly lower levels than wild-type MVs.

Although these data suggest that SpA could directly contribute to the IFN-ß response of macrophages to wild-type MVs, relatively little is known about the role of SpA in *S. aureus* MV formation. Therefore, further studies will be required to examine whether it is the presence of SpA itself in wild-type MVs that leads to the increased IFN-ß mRNA induction in macrophages, or if—consistent with the altered protein profile and protease protection of Δ*spa* MVs relative to wild-type MVs—deletion of SpA in *S. aureus* affects the biogenesis and/or cargo composition of MVs, thereby influencing the overall immunomodulatory activity of MVs produced by Spa-negative strains.

### A subset of immunomodulatory MV-associated nucleic acids are protected from nuclease degradation

Transfection reagents are known to facilitate RNA uptake by mammalian cells while protecting the RNA from nuclease degradation. Transfection of *S. aureus* RNA into innate immune cells was previously shown to trigger the production of Type I IFNs and pro-inflammatory cytokines^6–10^. We verified that RNA extracted from *S. aureus* whole-cells induce IFN-β mRNA expression in RAW 264.7 cells after 3h of transfection with lipofectamine (Fig. S7A). Notably, *S. aureus* RNA only triggered IFN-β production when it was delivered in complex with a tranfection reagent. This observation is in line with the hypothesis that immunomodulatory RNA derived from *S. aureus* can be transmitted into host cells via protective delivery vehicles, such as MVs.

We then set out to address the hypothesis that immunomodulatory RNA and DNA that are secreted in association with MVs are protected from nuclease attack. It is also possible that proteins on the surface of MVs may bind and protect MV-associated nucleic acids from nuclease activity. In an effort to determine the protection status and localization of RNA and DNA in relation to MVs, we conducted a series of nuclease protection assays. As visualized by native agarose gel electrophoresis, the results revealed dark bands in the untreated and RNase A-treated MV samples (Fig S8A; lanes 1 & 2) which were likely RNA and DNA that were unable to migrate from the sample loading wells into the agarose matrix due to their association with the large vesicles. However, we further detected MV-associated nucleic acids that were highly resistant to RNase A and DNAse I treatment even after MVs were subjected to combinations of heat, detergent, and Proteinase K digestion (Fig. S8A). We ruled out the possibility that components present in MV samples simply inactivated the nucleases by noting that exogenously added RNA or DNA mixed with MV samples could be degraded effectively by RNase A and DNase I, respectively (Fig. S4A & B).

Using RNAse A and DNase I to treat MVs yielded results that were surprising yet inconclusive: We expected MVs to protect associated nucleic acids, but it was unclear why a population of MV-associated nucleic acids remained protected from degradation even after we disrupted the vesicular structure. As such, we decided to employ benzonase, a nuclease which degrades all forms of RNA and DNA, to further examine this phenomenon. First, we verified that *S. aureus* RNA could be degraded by benzonase treatment (Fig. S7B). In addition to using agarose gels for these nuclease protection assays, we also assessed the MV-associated nucleic acids using denaturing urea polyacrylamide gel electrophoresis—which allows for more sensitive detection of smaller and low abundance RNA and DNA molecules compared to native agarose gels (Fig. 3). Nuclease protection assays on purified MVs revealed that a considerable population of nucleic acids are susceptible to degradation by benzonase digestion alone (Fig. 3 and Fig. S8B). Intriguingly however, neither disruption of the vesicle lipid bilayer by heating MVs to 95°C and/or treating with detergent or proteinase K lead to increased digestion of the benzonase-resistant nucleic acids. Altogether, these results suggest that a subpopulation of MV-associated nucleic acids is highly resistant to nuclease degradation even in conditions that disrupt the MV structure.

**Figure 3.**
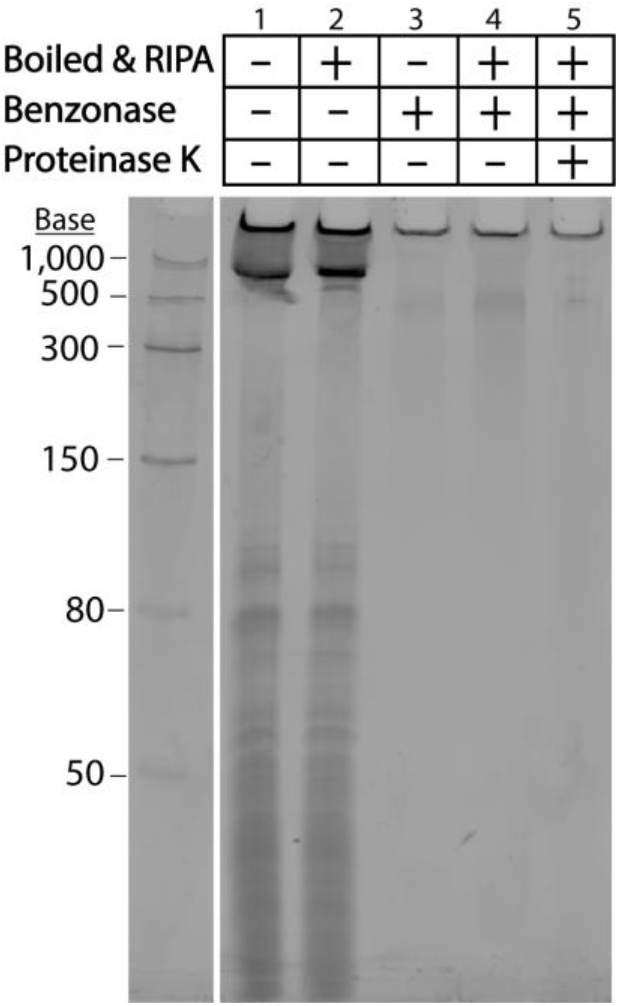
Detergent, Benzonase, and Proteinase K sensitivity of MV-associated nucleic acids. Nucleic acids from untreated MVs (10 μL, 7 μg total protein), or permeabilized MVs boiled at 95°C for 10 min followed by RIPA treatment for 10 min at 4°C, and/or MVs treated with Benzonase at 37°C for 30 min, and/or MVs subjected to Proteinase K digestion at 37°C for 30 min were separated on a 10% denaturing urea-PAGE gel in TBE buffer and stained with SYBR Gold. The full-length gel is presented in Supplementary Figure S12.

To characterize the MV-associated nucleic acids that were observed by gel electrophoresis, we extracted RNA and DNA from untreated MVs, benzonase-treated MVs (bMVs), or permeabilized + proteinase K treated bMVs (permeabilized/proteinased/bMVs), and we analyzed the specific RNA and DNA profiles using a Bioanalyzer. We found that the predominant RNA type that is secreted in association with untreated MVs is generally < 300 nt in length, while the predominant DNA species is ~500 base pairs long (Fig. 4A, B and Fig. 5A, B, respectively). Interestingly, benzonase treatment alone digested a significant portion of MV-associated RNA molecules, but permeabilization and proteinase K digestion of MVs prior to benzonase treatment did not significantly affect the amount of RNA extracted from MVs (Fig. 4C). Notably, the benzonase-susceptible RNA molecules included those that were ~200 nt long. A sharp, distinct RNA band slightly greater than 25 nt was resistant to benzonase treatment in all of the conditions tested, in accordance with our gel electrophoresis assays. We also noted that the 500 bp DNA band extracted from MVs was not affected by benzonase treatment alone or with additional MV permeabilization and proteinase K treatments (Fig. 5C). Collectively, these data along with the results from the various gel-electrophoresis-based assays provide support for the observation that subpopulations of benzonase-sensitive RNA molecules as well as subpopulations of benzonase-resistant RNA and DNA molecules are secreted in association with *S. aureus* MVs. It was furthermore particularly remarkable that detergent and proteinase K treatment did not lead to significant changes in the nuclease susceptibility of the MV-associated RNA and DNA.

**Figure 4.**
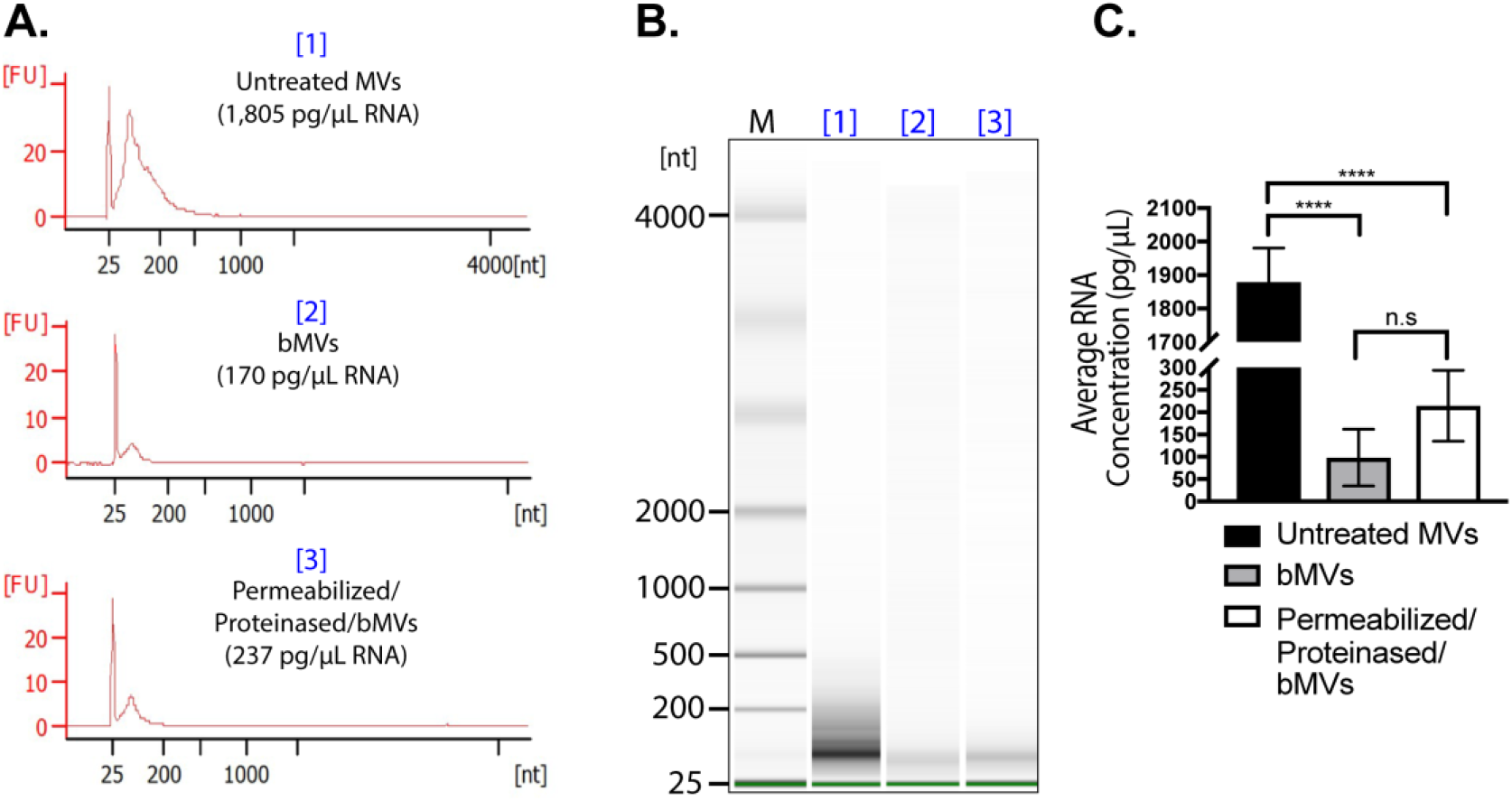
The RNA content of *S. aureus* MVs consists of Benzonase-sensitive and Benzonase-resistant sub-populations. (A) Bioanalyzer RNA profile of untreated MVs [1], RNA profile of MVs treated with Benzonase [2], and RNA profile of heated MVs treated with detergent, Proteinase K, and Benzonase [3]. X-axis represents the RNA nucleotide length and the y-axis represents fluorescent units. (B) Bioanalyzer gel image of RNA extracted from MV samples treated under different conditions. M indicates the relative migration of RNA standards. The complete gel image is presented in Supplementary Figure S13A. (C) RNA concentrations from each MV treatment were averaged from triplicate experiments. The data is represented as mean SEM from n = 3. Profiles are representative of two independent experiments. Statistical analysis was performed by one-way ANOVA with Tukey’s multiple comparison test (n.s = not significant; ****p < 0.0001).

**Figure 5.**
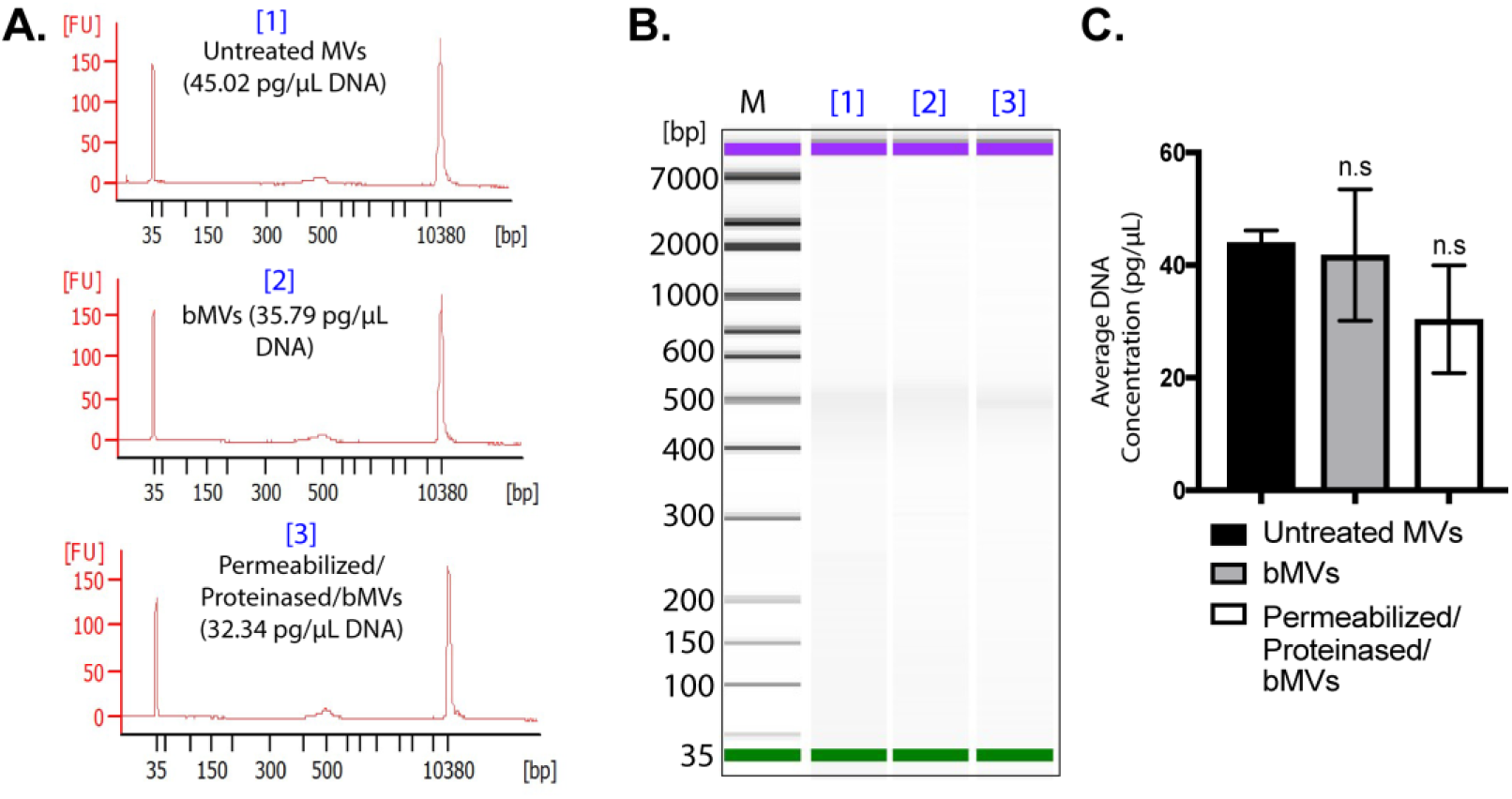
The DNA content of *S. aureus* MVs is resistant to Benzonase treatment. (A) Bioanalyzer DNA profile of untreated MVs [1], DNA profile of MVs treated with Benzonase [2], and DNA profile of heated MVs treated with detergent, Proteinase K, and Benzonase [3]. X-axis represents DNA base pairs and the y-axis represents fluorescent units. (B) Bioanalyzer gel image of DNA extracted from MV samples treated under different conditions. M indicates the relative migration of DNA standards. The complete gel image is presented in Supplementary Figure S13B. (C) DNA concentrations from each MV treatment were averaged from triplicate experiments. The data is represented as mean SEM, n = 3. Profiles are representative of two independent experiments. Statistical analysis was performed by one-way ANOVA with Tukey’s multiple comparison test (n.s = not significant).

With this characterization of MV-associated nucleic acids in mind, we next investigated the effect on IFN-ß induction of benzonase treatment of the MVs prior to their incubation with cultured macrophages (Fig. 6A). While IFN-ß mRNA expression was significantly reduced in response to bMVs relative to MVs, bMVs still induced potent IFN-ß mRNA expression in macrophages (Fig. 6B). These results demonstrate that IFN-ß mRNA is induced significantly by both benzonase-sensitive and benzonase-resistant subpopulations of MV-associated RNA and DNA molecules.

**Figure 6.**
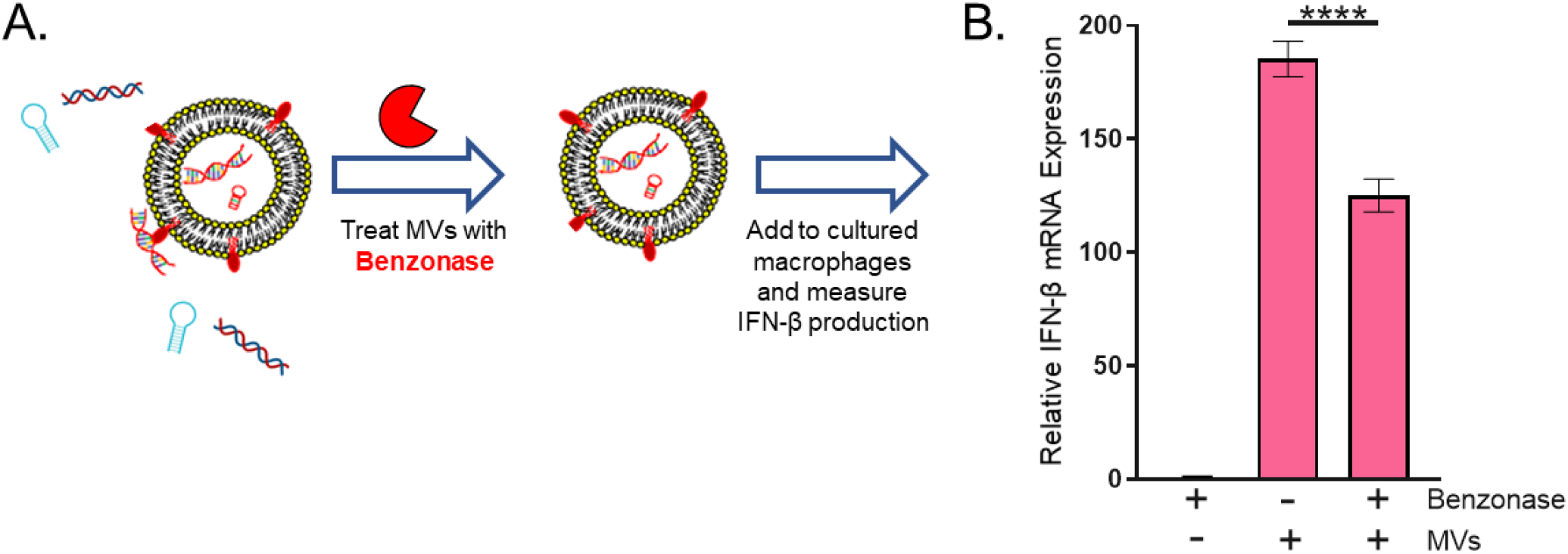
Benzonase-treatment reduces MV-mediated IFN-ß mRNA expression in macrophages. (A) Schematic representation of the experimental set-up. Purified MVs were treated with either Benzonase (250 U/μL) or HEPES-NaCl for 30 min at 37°C. A NT sample was prepared by incubating HEPES-NaCl with 1 μL Benzonase (250 U/μL) similarly to MVs. After 30 min, MVs were pelleted and re-suspended in HEPES-NaCl. Benzonase-treated and untreated MVs (5 μg/mL) were then incubated with RAW 264.7 cells for 3 h at 37°C in 5% CO_2_. (B) Macrophages were analyzed for expression of β-actin and IFN-β mRNA using qPCR. The data is represented as mean ± SEM from n = 3. Statistical analysis was performed by one-way ANOVA with Tukey’s multiple comparison test (****p < 0.0001).

### MVs induce IFN-ß mRNA expression largely through endosomal TLR signaling in macrophages

Generally, microbial nucleic acid recognition takes place in the cytoplasm or in endosomal compartments of host mammalian cells. In the case of *S. aureus*-derived nucleic acids, Type I IFNs can be produced via endosomal TLR signaling and through cytosolic Stimulator of Interferon Signaling Gene pathways^5,51^. Moreover, a recently published study demonstrated that *S. aureus* MVs are internalized into human macrophages primarily through dynamin-dependent endocytic pathways^35^. To test whether bMVs induce potent IFN-ß expression via endosomal- or cytoplasmic-dependent signaling, we treated macrophage cells with Dynasore, a dynamin-dependent endocytosis inhibitor, prior to stimulation with bMVs (Fig. 7A). Treatment with Dynasore significantly reduced the expression of IFN-ß mRNA in response to MVs, suggesting that entry of MVs and their immunomodulatory cargo could occur via dynamin-dependent endocytic pathways. It is well understood that endosomal acidification is required for TLR signaling activity^56^ (Fig. 7B). We further hypothesized that the IFN-ß mRNA produced in response to *S. aureus* MV nucleic acids have been released by acidification of the MV-containing endosomal compartment to activate cognate TLRs. To test this, we pretreated macrophages with the endosomal acidification inhibitors bafilomycin A1 and/or chloroquine^56^. Bafilomycin and chloroquine severely reduced IFN-ß mRNA expression in response to bMVs (Fig. 7C). To track the location of MV-associated RNA, MV-associated RNA was stained using the RNA-specific SYTO RNASelect-dye and incubated with the cultured macrophages for 10 min. Macrophages were subsequently fixed and immunofluorescently labeled for the early endosomal marker EEA1. Two fluorescent images were simultaneously acquired and analyzed for colocalization using STORM microscopy (Fig. 7D). To quantitate the degree of colocalization between RNA-stained MVs with EEA1 in macrophages, analysis of micrographs of the dual immunofluorescent labeling for MV-RNA and EEA1 was performed using the Mander’s overlap coefficient algorithm. Quantitative analysis revealed “moderate colocalization” between RNA-labeled MVs (green fluorescence) and the early endosomal marker EEA1 (red fluorescence) in macrophage cells. Because *S. aureus* MVs contain the IgG binding proteins Spa and Sbi^30,40,41^ (Fig. S5 C & D), we incorporated an irrelevant primary antibody control to account for potential non-specific binding of the EEA1 antibody to MV-associated Spa and Sbi in these immunofluorescence staining assays (Fig. S9). We show that moderate colocalization of bMV-associated RNA is observed in cells that were blocked both in the presence or absence of an irrelevant human IgG monoclonal antibody (Fig. S9A & B). Additionally, we provide evidence that the stained RNA is carried into RAW 264.7 cells in association with bMVs due to colocalization of DiD-stained bMVs with RNA-stained bMVs (Fig. S9C). These observations suggest that internalized MVs are capable of delivering RNA into endosomal compartments of host recipient cells.

**Figure 7.**
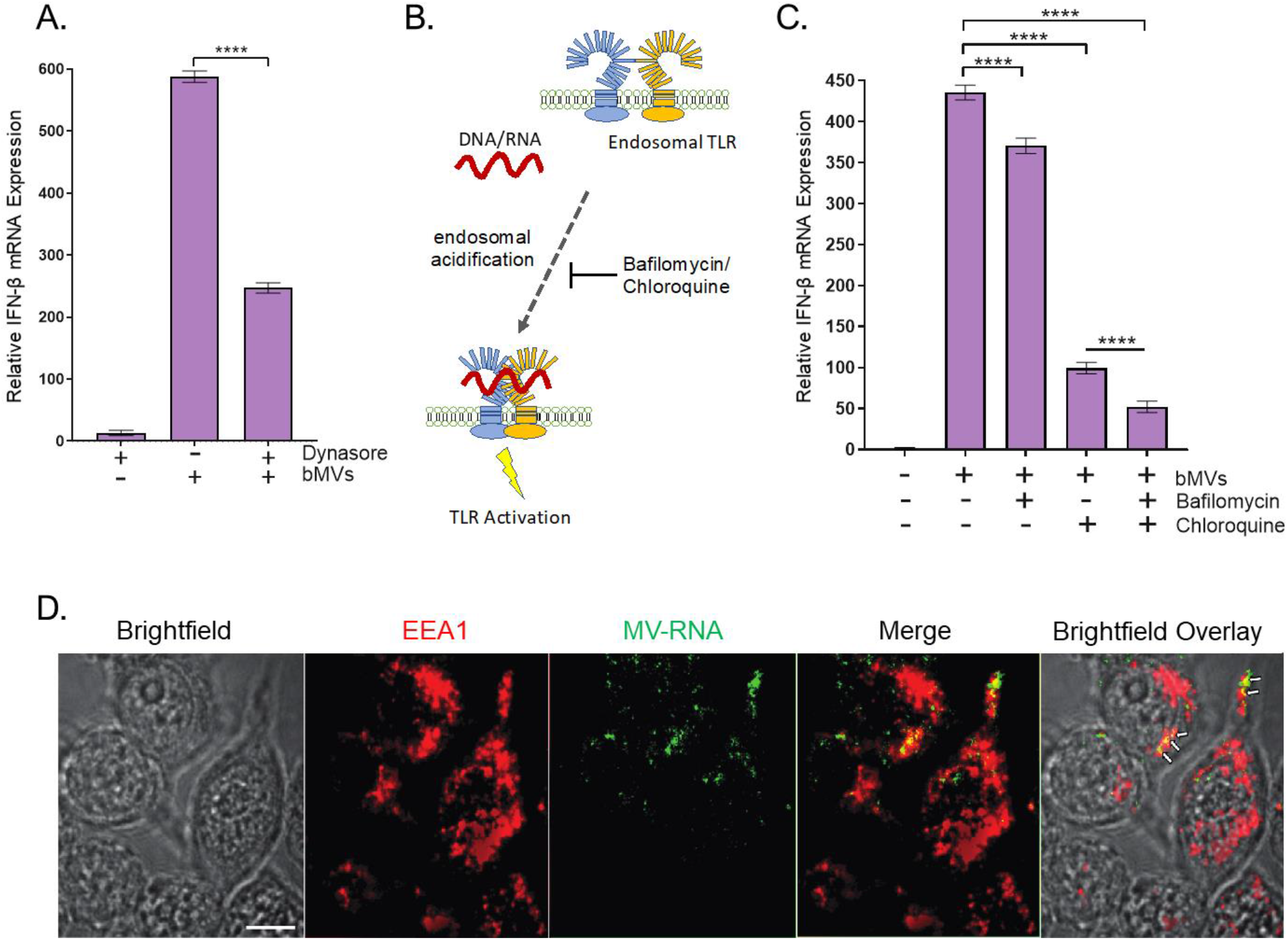
Dynamin-dependent endocytosis is likely involved in MV-mediated induction of IFN-β in RAW 264.7 cells. (A) RAW 264.7 cells were either treated with 80 μM Dynasore, which inhibits dynamin-mediated endocytosis, or with PBS buffer for 30 min and incubated with bMVs (5 μg/mL total protein) for 3 h, homogenized, and processed for qPCR analysis of β-actin and IFN-β mRNA expression. (B) Schematic representation of endosomal TLR signaling inhibition by the endosomal acidification inhibitors bafilomycin and chloroquine. Acidification of the endosome is essential for TLR3, TLR7, and TLR9 signaling in response to nucleic acid binding. Inhibition of the endosomal acidification process by bafilomycin A1 or chloroquine treatment prevents proteolytic cleavage of endosomal TLRs thereby inhibiting their signaling activity. (C) RAW 264.7 cells were either treated with PBS, 0.5 μM bafilomycin A1, and/or 100 μM chloroquine for 30 min followed by stimulation with bMVs (5 μg/mL) for 3h, homogenized, and analyzed by qPCR for β-actin and IFN-β mRNA expression. (D) Association of MV RNA with an early endosomal marker. Macrophages were incubated for 10 min with SYTO RNASelect-stained MVs (1 μg/mL), washed, fixed, and immunostained with anti-EEA1 antibodies. Arrows indicate “moderate colocalization” between MV-RNA and EEA1.

To develop a mechanistic framework for IFN-ß mRNA induction by *S. aureus* MVs, we examined the role of endosomal TLR signaling in the macrophage IFN-ß response to MVs. First, we sought to confirm that the bMV-mediated IFN-ß induction was dependent upon TLR signaling by abolishing all TLR-dependent signaling using MyD88/TRIF-deficient (*MyD88^-/-^Trif^-/-^*) BMDMs. MyD88 and TRIF are adaptor proteins essential for signaling downstream of TLR activation and are not required for cytoplasmic receptor signaling^57^. bMV treatment of *MyD88^-/-^ Trif^-/-^* macrophages resulted in significantly less IFN-ß expression compared to wild-type macrophages (Fig. 8A). This result confirms that TLRs are the dominant class of pattern recognition receptors (PRRs) that contribute to the IFN-ß signature response of macrophages to bMVs. Interferon regulatory factor 3 and 7 (IRF3 and IRF7) have been identified as critical regulators for nucleic acid-induced IFN production as they signal downstream of MyD88 and TRIF^58^. Since IRF3 and IRF7 can be activated by TLRs and by cytosolic nucleic acid receptors, we were interested in testing whether they are involved in the IFN-ß macrophage response to bMVs. Indeed, IFN-ß induction was also significantly reduced in *IRF3^-/-^/IRF7^-/-^* macrophages upon bMV stimulation compared to wild-type macrophages (Fig 8B). Combined, these results further support the conclusion that nucleic acids delivered by bMVs activate endosomal TLRs to induce a potent IFN-ß response in macrophages.

**Figure 8.**
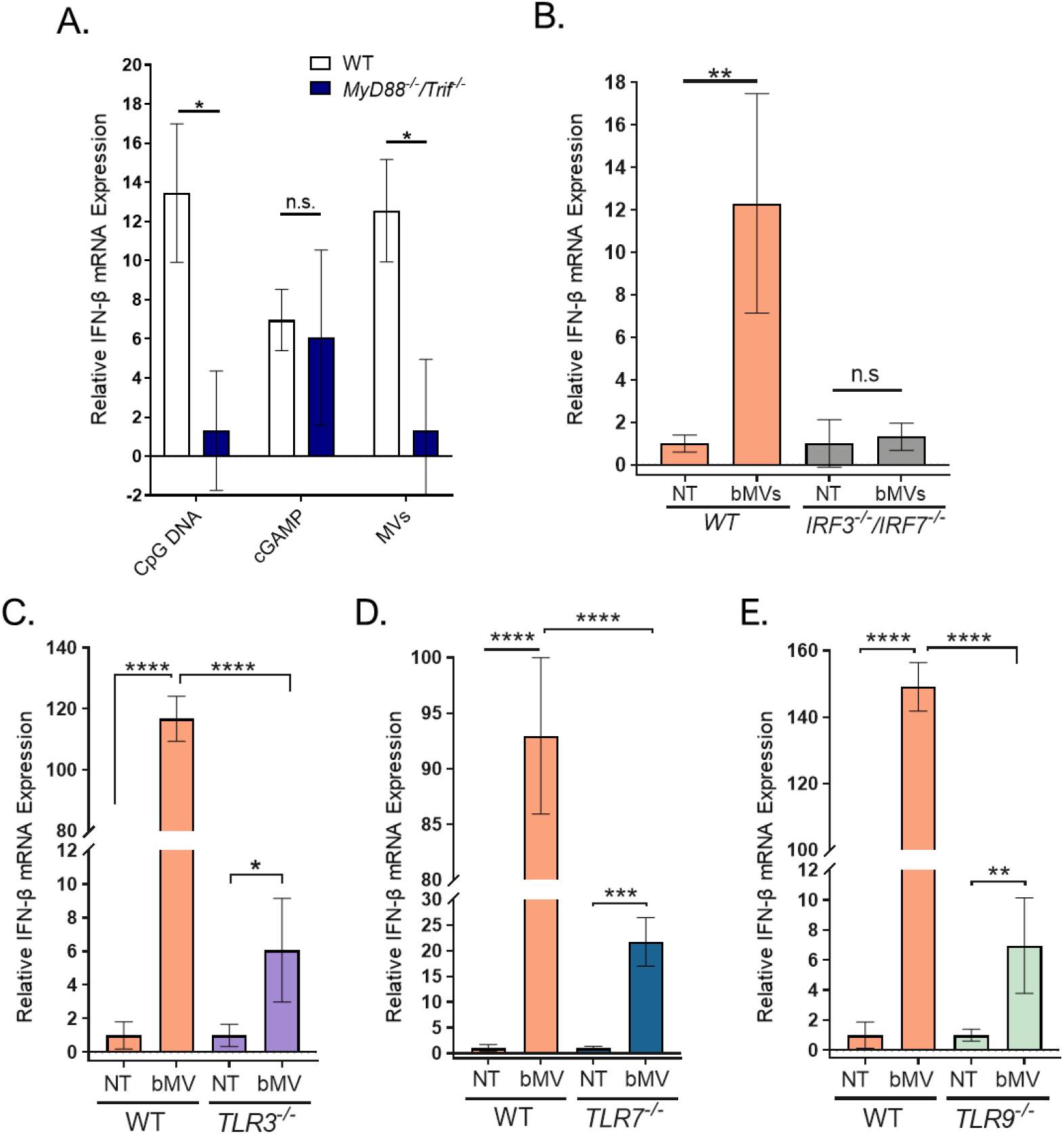
MV-associated RNA induces IFN-β largely through endosomal TLR signaling in murine macrophages. (A) Wild-type and *Myd88^-/-^/TRIF^-/-^* BMDMs were treated with 2 μM CpG DNA, 1 μg/mL 2’3-cGAMP, or bMVs (5 μg/mL total protein) for 3 h at 37°C, 5% CO_2_. (B-E) Wild-type, *IRF3^-/-^/IRFT7^-/-^, TLR3^-/-^, TLR7^-/-^*, *TLR9^-/-^* macrophage cells were stimulated with bMVs (10 μg/mL total protein) or with HEPES-NaCl (NT control) for 3 h. Treated macrophages were homogenized and β-actin and IFN-β mRNA expression levels were measured using qPCR. Data are expressed as means ± SE of triplicate samples and are representative of three independent experiments. n.s, not significant; * p ≤ 0.05; ** p < 0.005; *** p < 0.001; ****p < 0.0001.

Since bMV-mediated induction of IFN-ß expression was mostly dependent on endocytic TLR activation we investigated the role of TLR3, TLR7, and TLR9 in detecting bMV-associated RNA and DNA. First, we established that wild-type macrophages responded to purified bMVs in a dose-dependent manner (Fig. S10). Additionally, the functional properties of knock-out macrophage cells were verified by their reduced IFN-ß mRNA production in response to established IRF3/IRF7, TLR3, TLR7, and TLR9 agonists (Fig. S11). To explore the role of endosomal TLRs, we exposed *TLR3^-/-^, TLR7^-/-^,* and *TLR9^-/-^* macrophages to bMVs. Intriguingly, as evaluated by IFN-ß mRNA expression, the responsiveness of *TLR3^-/-^, TLR7^-/-^,* and *TLR9^-/-^* macrophages to bMVs was strongly impaired (Fig. 8C-E). Furthermore, we wondered whether TLR3 was also an important contributor to IFN-ß induction in RAW 264.7 cells. To explore this question, we pre-treated RAW 264.7 cells with a TLR3/dsRNA inhibitor prior to stimulation with either the TLR3 agonist Poly (I:C) (Fig. S12A) or with bMVs (Fig. S12B). As expected, the TLR3 inhibitor significantly reduced the IFN response of RAW 264.7 cells to Poly (I:C) (Fig. S12A). Interestingly, we also found that cells pretreated with TLR3 inhibitor produced a modest but statistically significant decreased level of IFN-β mRNA in response to bMVs (Fig. S12B). While TLR7 and TLR9 have previously been reported as sensors for *S. aureus-derived* nucleic acids^4,10^, our findings show for the first time a TLR3-dependent IFN-β induction by *S. aureus-* derived nucleic acids. Altogether, these data indicate that TLR3, TLR7, and TLR9 are all involved significantly in the signaling cascade that leads to the potent bMV-mediated IFN-β induction in cultured macrophages.

Next, we examined differences in the IFN-ß response to benzonased and untreated MVs in the various endosomal TLR-deficient macrophages to evaluate differences in signaling pathways induced by nuclease accessible and nuclease protected MV-associated nucleic acids. In accord with our data from RAW 264.7 cells, wild-type macrophages produced more IFN-ß mRNA after stimulation with untreated MVs compared to stimulation with bMVs (Fig. 9A). When *TLR3^-/-^* macrophages were stimulated with untreated MVs, IFN-ß mRNA expression levels were significantly increased compared to stimulation with bMVs (Fig. 9B). Intriguingly, benzonase treatment did not affect the ability of MVs to stimulate IFN-ß expression in *TLR7^-/-^* or *TLR9^-/-^* macrophages (Fig. 9C and 9D). These data suggest that a substantial portion of the stimulatory RNA and/or DNA content of *S. aureus* MVs is protected and can be delivered into host cells by MVs to activate intracellular nucleic acid receptor, but also that a population of exposed MV-associated RNA and/or DNA that enter into macrophages can signal through intracellular receptors that include TLR7 and TLR9.

**Figure 9.**
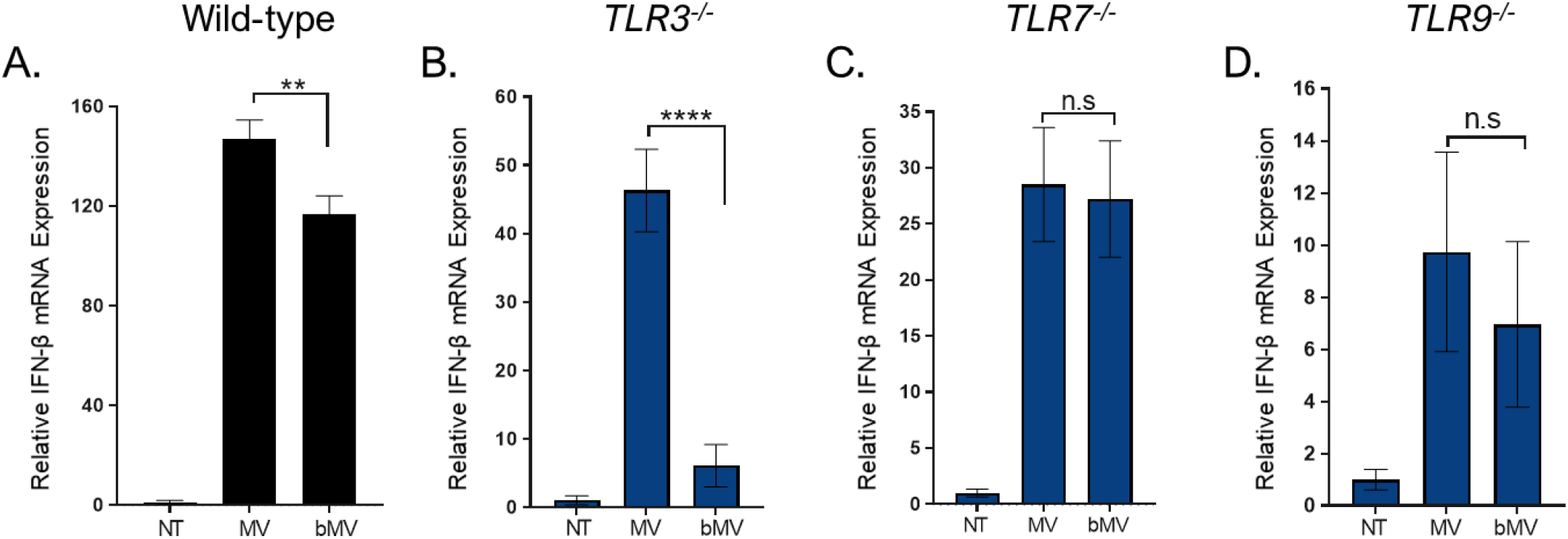
Treating MVs with benzonase reduces the IFN-β mRNA expression in both WT & *TLR3^-/-^* macrophages compared to macrophages stimulated with untreated MVs. (A-D) Wild-type, *TLR3^-/-^TLR7^-/-^, TLR9^-/-^* macrophage cells were stimulated with untreated MVs (10 μg/mL total protein), bMVs (10 μg/mL total protein), or with HEPES-NaCl (NT) for 3 h. Treated macrophages were homogenized and β-actin and IFN-β mRNA expression levels were measured using qPCR. Data are expressed as means ± SE of triplicate samples and are representative of three independent experiments. n.s, not significant; **p < 0.005; ****p < 0.0001.

Lastly, due to what appears to be a significant contribution by RNA to the potent bMV-mediated IFN-ß induction in macrophages, we asked whether nuclease-protected RNA molecules could be delivered into wild-type macrophages via bMVs. Based on the uptake of SYTO RNASelect-labeled MVs into RAW 264.7 cells shown in Figure 7D, we investigated the delivery of bMV-associated RNA into wild-type macrophages by staining bMV-RNA with SYTO RNASelect. Furthermore, we also sought to confirm that the bMV-RNA detected in macrophages is specifically of staphylococcal origin by metabolically labeling *S. aureus* RNA with the nucleoside analog of uracil, 5-ethynyl uridine (5-EU). We first performed a proof-of-concept experiment to demonstrate that the Click-iT RNA Alexa Fluor 488 Imaging Kit could be adapted to successfully label the RNA of actively proliferating *S. aureus* cells in culture (Fig. S13). To achieve this, we supplemented the culture media (RNased NB) with 100 μM 5-EU and grew *S. aureus* cells to mid-log phase, which allowed 5-EU to become incorporated into newly synthesized RNA. This technique effectively labeled 5-EU RNA in *S. aureus* cells grown to mid-log phase (Fig. S13B).

To examine uptake of bMVs and their RNA cargo, we scaled up the volume in which 5-EU labeled *S. aureus* cells were grown, we supplemented the culture media with 150 μM 5-EU, and isolated MVs from the supernatant of the culture after reaching mid-log phase. Purified MVs were treated with benzonase and the MV lipids were labeled with DiD, a lipophilic dye. We also purified MVs from *S. aureus* cells grown in RNased NB without 5-EU, we treated the MVs with benzonase, labeled their membranes with DiD and the associated RNA molecules with SYTO RNASelect. Labeled bMVs were incubated with wild-type macrophages for 1h, the macrophages were washed to remove unbound or non-internalized bMVs, followed by fixation and permeabilization. 5-EU RNA bMVs were labeled within macrophages using the Click-iT RNA labeling kit. DiD-bMVs and 5-EU RNA bMVs were readily taken up by wild-type macrophages after 1h of co-incubation (Fig. 10A and Fig. S14A & B). We also show that DiD + SYTO RNA bMVs were internalized by macrophages in a similar fashion (Fig. 10B and Fig. S14C). These findings are consistent with the suggestion that bMVs and their nuclease-protected, bacterially-derived RNA cargo are internalized into wild-type macrophages.

**Figure 10.**
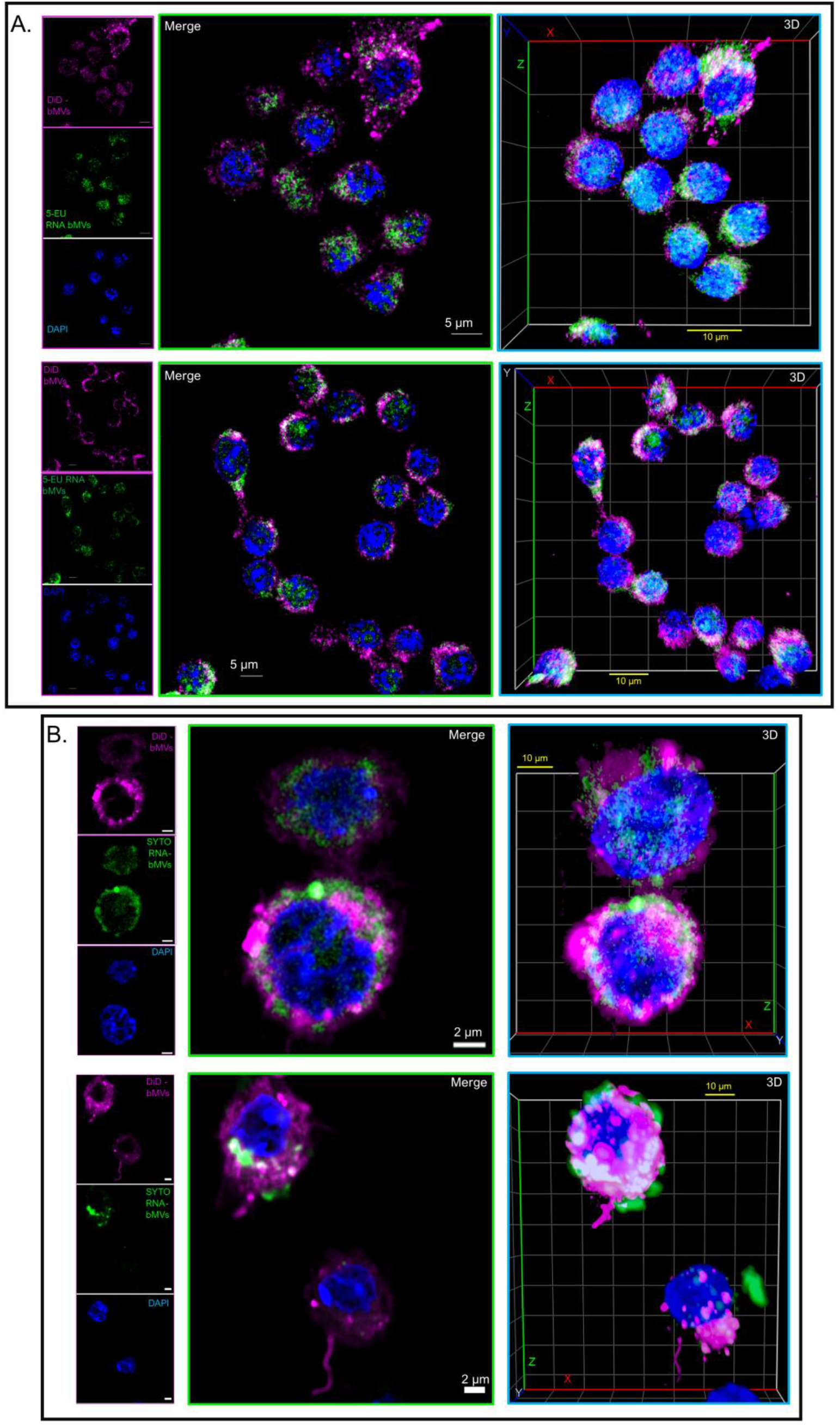
*S. aureus* bMVs and their associated RNA cargo is delivered into wild-type macrophages. (A) Wild-type macrophages were treated with DiD-labeled bMVs (magenta) for 1h. Macrophages were then fixed and permeabilized using saponin. *S. aureus* 5-EU RNA in macrophages was reacted and stained with the Click-iT Alexa Fluor 488 microscopy labeling kit (green), and counterstained with DAPI (nucleus; blue). Representative z-stack projections and their corresponding 3D reconstructions are shown for each of the 2 independent experiments that were performed. N = 64 stacks (top panel) and N = 64 stacks (bottom panel). (B) WT macrophages were treated with DiD + SYTO RNA-labeled bMVs (green; bMV RNA & magenta; bMV lipids) for 1h. Macrophages were then fixed, permeabilized, and stained with DAPI. Representative z-stack projections and their corresponding 3D reconstructions are shown for each of the 2 independent experiments that were performed. N = 40 stacks (top panel) and N = 64 stacks (bottom panel). Images demonstrate that bMV-associated RNA molecules can be delivered to macrophages and possibly released into the cytoplasm and peri-nuclear region after 1 h of co-incubation. All images were captured using a Zeiss LSM 880 equipped with Airyscan (63x objective). Scale bars are indicated.

## Discussion

Microbial-derived RNA has recently received attention as a PAMP capable of modulating host immune responses via activation of signaling pathways previously considered to respond primarily to viral RNA^1–3^. In light of these reports, both RNA and DNA of bacterial origin have now been described as strong inducers of Type I IFNs by activating cytosolic and/or endosomal receptors. That bacterial RNA and DNA can drive host immune responses and determining whether such responses are protective or detrimental to the host has been the subject of the majority of studies on immunomodulatory nucleic acids^1–3,5^. In comparison, fewer studies have examined how those immunomodulatory nucleic acids are secreted by bacteria in general–and by the extracellular human pathogen *S. aureus* in particular—and how they are transferred to host cells to eventually reach IFN signaling receptors^2,3,18–20^. In conjunction with an increasing number of studies showing secretion of bacterial RNA and DNA via extracellular vesicle production^18–20^, we hypothesized that *S. aureus* secrete MVs containing bacterially-derived RNA and DNA molecules that can withstand nuclease degradation and be delivered into host cells. In this report we show for the first time that in cultured mouse macrophages, IFN-ß mRNA induction can be triggered by purified *S. aureus* MVs. We further provide evidence that MV-induced IFN-ß expression requires endosomal TLR activation and is specifically promoted by engagement of endosomal TLR3, TLR7, and TLR9 receptors—pointing to the likelihood that MV-associated, protected RNA and DNA molecules are inducing such a response.

Nuclease protection assays revealed subpopulations of nucleic acids that are protected from nuclease degradation. This finding was consistent with several previous studies on OMV-associated nucleic acids which suggested that OMVs provide a physical barrier that blocks extracellular nucleases from accessing their associated RNA and DNA cargo. Surprisingly, even when the structure of MVs was challenged with detergent and proteinase K treatments, we found that certain RNA molecules were resistant to nuclease degradation. Nuclease protection assays of *P. aeruginosa* OMVs by Koeppen et al. indicated that all OMV-associated RNA species were protected from RNase degradation^12^. OMVs from *Salmonella enterica spp. Typhimurium* were found to be protective for a selection of non-coding RNAs^59^. Additionally, RNase A treatment on *Vibrio cholerae* and *Escherichia coli* OMVs were found to be inefficient for complete removal of OMV-associated RNA^15,60^. These observations of RNA protection may be due to the presence of secondary structures and/or non-proteinaceous binding partners that protect the nuclease-resistant RNA from degradation.

Interestingly, we found that all of the detectable MV-associated DNA was resistant to benzonase treatment in all of the conditions tested. However, because there were no significant changes in the amount of DNA extracted from MVs that underwent membrane damage, we were unable to determine the exact nature of the association between DNA and MVs. Our data contrast reports concluding the external association of nucleic acids with OMVs in cases where DNase treatment alone was sufficient to degrade the DNA (plasmid and/or chromosomal fragments) associated with OMVs secreted by Gram-negative pathogens^16,17^. In other Gram-negative species such as *S. Typhimurium, Porphyromonas gingivalis, and P. aeruginosa,* chromosomal DNA fragments were determined to be both external and internal to OMVs^16,17,61^. Collectively, the inconsistent reports on vesicle-associated RNA and DNA suggest that, depending on the microbe, vesicles differ in their RNA and DNA cargo composition and location with respect to the outermost vesicle membrane. These variations may also be attributed to different vesicle biogenesis mechanisms, which would influence the surface features and localization of lumenal cargo into vesicles.

Wang *et al.* demonstrated that inhibition of dynamin-dependent endocytosis prevented internalization of *S. aureus* MVs into human macrophages^35^. We observed that IFN-β mRNA expression is significantly reduced in dynasore-treated macrophages. Thus, dynamin-dependent endocytosis is strongly implicated in the MV-mediated induction of IFN-β mRNA in macrophages. However, based on these data alone we are unable to ascertain the extent to which this reduced IFN-β response was due to the blockage of MV uptake through dynamin-dependent endocytic pathways or if this was due to other factors. In fact, upon stimulation with their respective ligands, TLR3 and TLR9 are translocated from the endoplasmic reticulum to the cell surface before being internalized through dynamin-dependent endocytosis into endosomal compartments where they initiate signaling events^62–64^. Therefore, it is possible that dynasore-treated macrophages have a diminished IFN-β response due to the combined inability of certain MVs from being internalized through endocytosis and the inability of TLR3 and TLR9 to localize into endosomal compartments, which could prevent effective sensing and activation of MV nucleic acids by these TLRs.

Internalization of vesicle-associated DNA and RNA into host cells has been demonstrated recently for OMVs. OMVs derived from uropathogenic *E. coli* were reported to deliver 5-EU RNA to the nucleus of human bladder cells after 15 h of incubation^15^. Metabolically labeled DNA from *P. aeruginosa* OMVs localized to the nucleus of A549 epithelial cells after 5 h of treatment and intracellular trafficking assays indicated that OMV-DNA is transported to the nucleus over the course of 18 h^17^. By metabolically labeling the RNA cargo of bMVs, we provide visual evidence that bMV-associated RNA is delivered into macrophages after 1 h of incubation. We also showed that certain populations of MV-associated RNA colocalized to early endosomal compartments in RAW 264.7 cells in as early as 10 min. Importantly, the work presented here showed delivery of bMVs and their stably associated RNA cargo into macrophages during time points leading up to the IFN-β peak induction of 3 h. Although follow-up investigations will be required to assess specific internalization pathways and trafficking of MV-associated nucleic acids in host immune cells, the work presented here suggests that certain populations of MV-associated RNAs could be trafficked into locations within cells where intracellular RNA receptors reside.

We also investigated the mechanism by which IFN-β mRNA production was stimulated by the internalized *S. aureus* MVs by blocking endocytic processing. Previous studies of MV-mediated signaling indicated that rupturing of the MV lipid bilayer facilitates targeted and delayed release of their macromolecular cargo to target cells, leading to a concentrated delivery of bioactive agents^65,66^. Bielaszewska *et al.* reported that a pore-forming toxin associated with enterohemorrhagic *E. coli* OMVs required acidic pH conditions to dissociate from OMVs to induce apoptosis^65^. We found that endosomal acidification also plays a role in the *S. aureus* bMV-induced IFN-β response in macrophages, as signaling was significantly reduced by bafilomycin and chloroquine treatments. This effect likely highlights a requirement for MVs to be processed within acidic endosomal compartments in order to release their immunomodulatory cargo for subsequent recognition by TLRs. A deeper understanding of how and when the immunomodulatory cargo is released from MVs and consequently interacts with cognate TLRs will require further investigation.

On the basis of our observations that lipofectamine-assisted delivery of *S. aureus* RNA is necessary for induction of IFN-β m RNA expression in RAW 264.7 cells, and that transfected Poly (I:C) elicited significantly higher IFN-β mRNA expression in wild-type and *TLR3^-/-^* cells compared to when it was delivered alone (Fig. S11B and C), we postulated that MVs play a crucial role as transport vehicles in the inter-kingdom communication of immunomodulatory RNAs. We also noted a significant increase in IFN-β induction by transfection of CpG DNA in wild-type cells (Fig. S11E) which could signify that the uptake and immunomodulatory activity of DNA molecules is enhanced upon complexation with lipid-based transfection reagents.

We investigated the hypothesis that MV-associated RNA and DNA molecules are immunomodulatory by assessing the contribution of endosomal nucleic acids receptors to the IFN-β response. To develop an understanding of the downstream specific signaling pathways that contribute to the MV-mediated IFN-β mRNA response in macrophages, we examined the IFN-β response in WT cells compared to the response that remains when each endosomal TLR receptor is eliminated. It should be noted that this general approach has limitations related to the possibility that multiple pathways could each make small contributions to the activation of IFN-β mRNA. Nevertheless, these studies allow for the identification of dominant PRR sensors of MV-associated cargo, thereby representing an important first step towards developing a full understanding of the MV-mediated delivery of immunomodulatory nucleic acids into host cells. In finding that, relative to WT macrophages, the strongly induced IFN-β mRNA expression is significantly diminished in macrophages lacking Myd88 and TRIF, IRF3 and IRF7, TLR3, TLR7 or TLR9, the data highlight the dominant roles of endosomal TLR pathways in the Type I IFN response to *S*. *aureus*-derived MVs. The murine-specific endosomal receptor TLR13 and its functional human homolog TLR8 are additional innate immune receptors that have previously been shown to initiate signaling cascades in response to *S. aureus* RNA^6–9^. Because we did not test TLR13 or TLR8-deficient cells in this study, we cannot rule out the possibility that TLR13 and/or TLR8 could also recognize MV-associated RNA and contribute to IFN-β signaling in host cells. Consistent with our findings, OMVs secreted by the respiratory pathogen, *Moraxella catarrhalis,* induced T-independent B cell activation partly by signaling through the endosomal DNA receptor TLR9^16^. Additionally, OMVs produced by the periodontal pathogens *P. gingivalis* and *Tannerella forsythia* were found to moderately stimulate TLR7, TLR8, and TLR9 as assayed in HEK-Blue cell lines^67^. Therefore, these endosomal receptors are utilized to recognize nucleic acids associated with both Gram-positive and Gram-negative bacteria-derived vesicles.

From the total stimulatory RNA and DNA that is associated with MVs, certain subpopulations are protected from benzonase degradation and those protected nucleic acids can be delivered into host cells and activate TLR3, TLR7, and TLR9. We speculate that another subpopulation of RNA and DNA molecules associate with MVs in a manner that leaves them susceptible to nuclease degradation, and these susceptible nucleic acids (in the absence of nucleases) can be delivered via MVs into host cells to stimulate IFN-β expression, likely through TLR7 and TLR9 activation. Although we detected a prominent subpopulation of RNA molecules that can be degraded by benzonase treatment alone, we did not detect benzonase-susceptible DNAs in purified MVs. Therefore, it is possible that additional subpopulations of MV-associated immunomodulatory DNA were present in MV samples but that their relative amounts were below the limit of detection for the assays used herein.

The signaling of *M. catarrhalis* OMV-associated DNA in B cells was previously shown to be abolished with DNase treatment of OMVs^16^. These observations indicated that surface-localized DNA was the predominant form of the vesicle-associated active nucleic acid. In a different study, DNA molecules that were associated internally with *P. aeruginosa* OMVs were compared to the types of DNA molecules that were external to the OMVs^17^. These authors hypothesized that DNA encapsulated by OMVs may play a different role in bacterial cell-cell communication as compared to DNA associated to the surface of OMVs. This hypothesis parallels differences observed for toxin delivery, in that soluble *S. aureus* α-hemolysin was reported to be less cytotoxic to HaCaT keratinocytes by inducing apoptosis-like cell death, whereas MV-enclosed α-hemolysin caused necrosis and atopic dermatitis-like skin inflammation in mice^68^. The data presented in this study suggest that both unprotected and protected *S. aureus* MV-associated nucleic acids induce significant responses in host cells, but do so by different pathways, and these differences could have as-yet-unexplored consequences. Future studies will focus on characterizing the immunomodulatory RNA and DNA molecules that associate with MVs but are susceptible to nuclease degradation.

Altogether, our data support the model that *S. aureus* MVs are internalized by macrophages and that they are likely trafficked to endosomal compartments to elicit a potent IFN-β expression via TLR signaling (Fig. 11). To our knowledge, this report provides the first piece of evidence that *S. aureus* releases MVs that are internalized by mouse macrophages to induce IFN-β via endosomal TLR-dependent signaling. In bacterial infections, type I IFNs can act as activators of protective immune responses or mediate immunosuppressive functions leading to a reduced ability of the host to clear bacterial infections^1–3,5,69,70^. The ambivalent role of type I IFN during bacterial infections is currently the topic of multiple investigations. Future studies will be necessary to address whether the potent IFN-β induction of *S. aureus* MVs has an impact on the pathogenesis of staphylococcal disease.

**Figure 11.**
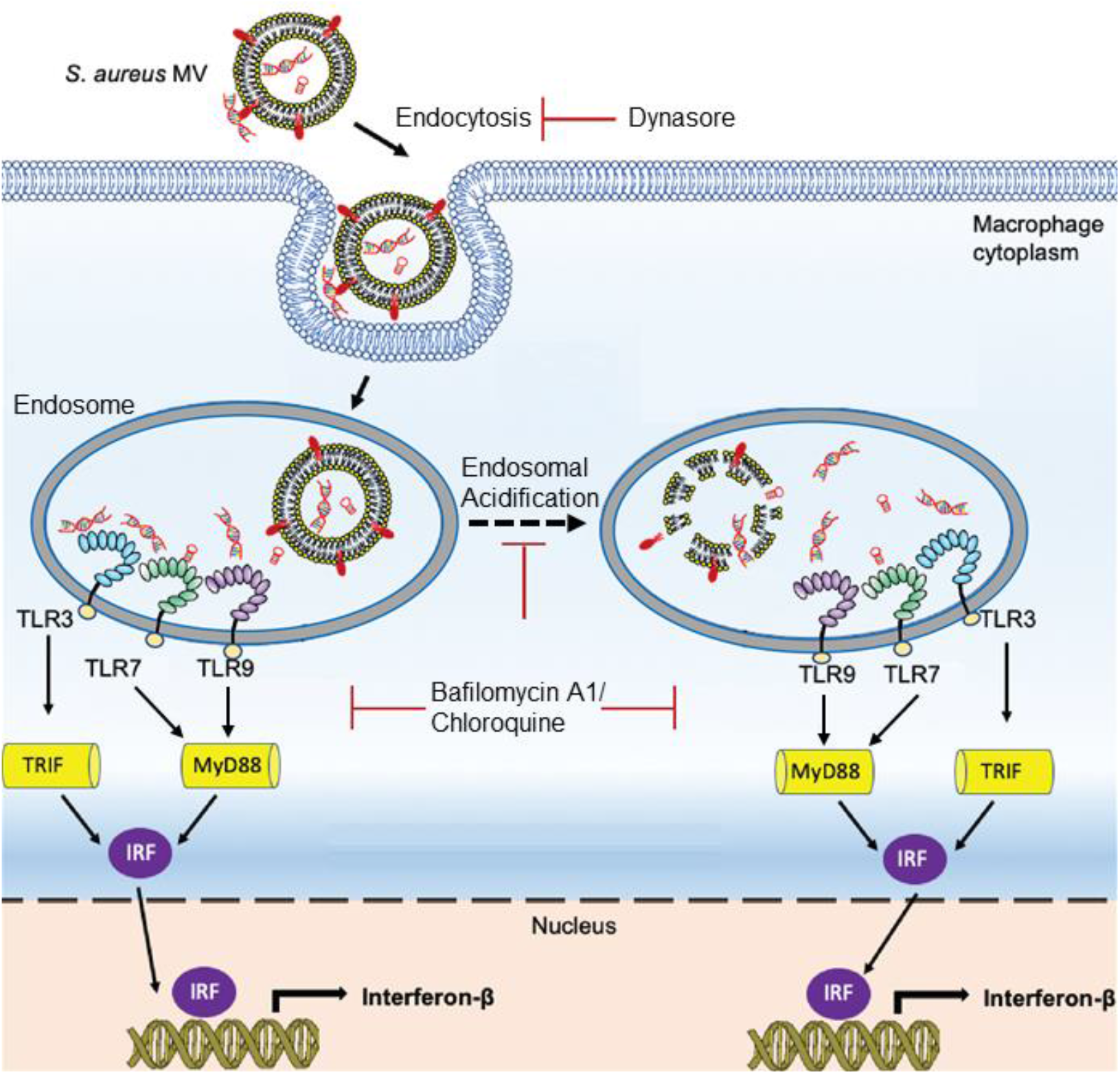
Proposed model for the MV-mediated delivery and trafficking of immunomodulatory RNA and DNA molecules into macrophages. *S. aureus* MVs carrying protected and non-protected nucleic acids are internalized by macrophage cells and MV-associated RNA and DNA are subsequently trafficked to endosomal compartments. After endosomal acidification, MV-associated RNA and DNA molecules are either separated from MVs to access TLR binding sites and/or non-protected DNA and RNA bind to PRRs through yet-to-be-determined mechanisms. MV-associated nucleic acids induce a potent IFN-β innate immune response by activating endosomal TLR signaling pathways. Downstream TLR signaling requires Myd88, Trif, IRF3, and IRF7 activation.

## Methods

### Bacterial strains and growth conditions

*S. aureus* Newman and Newman Δ*spa* were grown in nutrient broth (NB; 0.5% peptone, 0.3% beef extract, pH 7.0) overnight at 37°C with shaking (200 rpm). For MV nucleic acid and macrophage stimulation assays, *S. aureus* Newman wild-type and Newman Δ*spa* were grown in NB treated with 0.5 μg/mL RNase A (Thermo Fisher) at 37°C for 1.5 h prior to sterilization by autoclaving. Overnight cultures were sub-cultured to a 1/1000 dilution and grown for 4-5 hours at 37°C to an optical density of 1.0 at 600 nm (1.0 OD_600_).

### Bacterial growth curve and membrane integrity assay

Bacterial growth curves were recorded using a Spectramax 190 plate reader from Molecular Devices (California, US). A 96-well, U bottom plate was loaded with NB controls and with overnight *S. aureus* cultures diluted to 0.1 OD_600_ in fresh NB. The assay plate was incubated at 37°C for 18 h with constant shaking and the OD_600_ recorded every 15 min. Newman wild-type growth curve experiments were carried out in triplicate and representative growth profiles are shown.

For membrane integrity assays using the membrane-impermeant nucleic-acid dye SYTOX Green, *S. aureus* was grown to mid-log phase (1.0 OD_600_; 5 x 10^9^ cfu ml^-1^) and 1 mL culture samples were centrifuged at 10,000 x g for 10 min. The supernatants were discarded, and the pellets were resuspended in 10 mM HEPES, 150 mM NaCl, pH 7.5 (HEPES-NaCl). One of the resuspended cell pellets was heat-killed (80°C, 1 h) and used as a positive control. HEPES-NaCl buffer alone was used as a negative control. The samples were incubated (10 min, 20°C, in the dark) with 1 μM (final concentration) of SYTOX Green (Molecular Probes). SYTOX Green binding to bacterial DNA was measured using a microplate reader (Molecular Devices SpectraMAX GeminiXS) using 500 nm excitation /550 nm emission. Relative fluorescence units (RFU) were subtracted from the negative control RFU for each sample.

Live and dead *S. aureus* cultures were prepared according to the manufacturer’s instructions provided with the BacLight Kit (Invitrogen). Two flasks of 25 mL *S. aureus* cultures were grown to early-stationary phase (1.6 OD_600_). Bacterial cells were harvested using centrifugation at 10,000 *x* g for 10 min. After discarding the supernatant, each pellet was resuspended in 2 mL Tris Buffered Saline (TBS). One mL of this suspension was added to either 20 mL of TBS (for live bacteria) or to 20 mL of 70% isopropyl alcohol (for killed bacteria). Samples were incubated at room temperature for 1 h and the cells were pelleted by centrifugation at 10,000 x g for 15 min. Pellets were washed in TBS twice and ultimately resuspended in 10 mL TBS. For cell staining, working solutions of propidium iodide (PI; 20 mM in DMSO) and SYTO9 (3.34 mM in DMSO) were established by combining equal volumes of each stain and mixed thoroughly. 3 μL of the dye mixture was added to 1 mL of bacterial suspension, mixtures were incubated at room temperature (RT) in the dark for 15 min and 5 μL of the stained bacterial suspension was trapped between a slide containing a drop of BacLight mounting oil and a poly-L-lysine coated coverslip (#1.5). Confocal laser scanning microscopy (CSLM) was carried out using a Zeiss LSM 880 inverted confocal microscope and images were acquired using a 40x/1.2 water immersion objective (Duke Light Microscopy Core Facility). The 488 nm laser was used to acquire SYTO9 signals and the 594 nm laser was used to acquire signals from PI-stained cells. Images were processed using the ZEN 2.3 software (Carl Zeiss) and PI and SYTO9-stained cell counts were obtained using the Analyze Particles function available in ImageJ.

### Membrane vesicle isolation

MVs were isolated by adapting a previously described protocol^39^. Briefly, 6 liters of *S. aureus* Newman were grown in four 3-liter flasks to mid-exponential phase (OD_600_ of 1.0) and the culture was centrifuged (10,000 x g, 20 min, 4°C). The cell-free supernatant was filtered through a 0.45 μm pore size, hydrophilic polyvinylidene fluoride filter (Merck Millipore Germany). Filtered supernatants were concentrated 24-fold by ultrafiltration using a tangential flow system with 100-kDa hollow-fiber membrane (Pall Corporation) followed by ultracentrifugation (100,000 *x* g, 3 h, 4°C) to separate MVs from cell debris and contaminants. Pelleted vesicles were resuspended in HEPES-NaCl and subjected to an additional ultracentrifugation step (200,000 x g, 1 h, 4°C). MV preparations were tested for sterility by streaking a portion of the sample on NB agar and incubating at 37°C for 18 h.

### Density gradient centrifugation

Crude MV samples were adjusted to 50% (vol/vol) Optiprep (Sigma) solution and loaded to the bottom of a 13.2 ml ultracentrifuge tube. The sample was overlaid with a 40% Optiprep layer followed by a 10% Optiprep layer. The tube was centrifuged at 130,000 x g (Beckman SW 41Ti Swinging-bucket rotor) for 16-18 hours at 4°C and 2-ml fractions were collected from the top of the density gradient. The Optiprep solution was then removed by washing each fraction with HEPES-NaCl buffer in Oakridge tubes (Type 50.2 Ti fixed-angle rotor, 150,000 × g, 1 h, 4°C). Each fraction was resuspended in HEPES-NaCl buffer and 10 μL of each fraction was examined for protein concentration and lipid content using a Bradford protein assay (VWR) and the lipophilic FM4-64 dye (Molecular Probes), respectively. MV preparations were used in experiments according to their total protein concentration. Each fraction was also examined for DNA content by staining 10 μL of each fraction with SYTOX Green dye (1 μM final concentration, 500 nm excitation/550 nm emission) and MV RNA was determined by using the membrane permeable SYTO RNASelect green fluorescent cell stain (Invitrogen, 1 μM final concentration) to stain 10 μL of each fraction. SYTO RNASelect binding to MV RNA was measured (490 nm excitation/550 nm emission). MV fractions containing high proteolipid and nucleic acid content (F1-F3) were combined, checked for sterility, and stored at −20°C.

### Transmission electron microscopy imaging

10 μL of purified MV fractions were placed on 400-mesh copper grids (Electron Microscope Sciences) and stained with 2% uranyl acetate. All grids were viewed in a FEI Tecnai G^2^ Twin electron microscope at an 80-kV accelerating voltage and images were obtained using an AMT XR-60 charge-coupled device digital camera system. To quantitate diameters, MVs on the micrographs were measured from the outermost points of the stained membrane with the measurement function in the NIH FIJI Software.

### Cell culture strains and growth conditions

RAW 264.7 macrophage cells (ATCC TIB-71), macrophage cells derived from wild-type mice (NR-9456), TLR3, TLR7, TLR9 knockout mice, and from IRF3/7 double knockout mice (BEI Resources, NIAID, NIH) were maintained in Dulbecco’s Modified Eagle Medium (DMEM) containing high glucose (4.5 g/L), L-glutamine (4.5 g/L), and sodium pyruvate (110 mg/L). The growth medium was supplemented with 10% (v/v) heat-inactivated fetal bovine serum (Gibco). Wild-type (WT) and *MyD88^-/-^ TRIF^-/-^* bone marrow-derived macrophages (BMDMs) were cultured in RPMI 1640 + 20% (vol/vol) FBS + 12% (vol/vol) macrophage colony stimulating factor-conditioned media. All cell lines were grown under a humidified 5% CO_2_ atmosphere at 37°C.

### S. aureus RNA extraction

*S. aureus* was grown overnight in NB medium at 37°C, with shaking. The following day, the cells were diluted 1:50 in 5 mL fresh NB. Cells were harvested after reaching OD_600_ of 1 by centrifugation at 14,000 x g for 2 minutes. The pellet was resuspended in 200 μL lysozyme from a 3 mg/mL stock solution and 10 μL lysostaphin from a 10 mg/mL stock solution (Sigma-Aldrich). Cells were incubated at 37°C for 15 min. Total RNA was then extracted using the SV Total RNA Isolation System (Promega) according to the manufacturer’s instructions.

### SDS-PAGE and LPS staining

Newman wild-type and Δ*spa* MVs of equal amounts (2 μg total protein) and LPS standard from *E. coli* serotype 055:B5 (250 μg/mL) were mixed with 0.5x volumes of Laemmli sample buffer (65.8 mM Tris-HCL, pH 6.8, 26.3% w/v glycerol, 2.1% SDS, and 0.01% bromophenol blue; Bio-Rad) and boiled for 5 min prior to loading on 4-20% SDS-PAGE gradient gels (Mini-PROTEAN TGX precast gels; Bio-Rad). Precision Plus Protein Unstained Protein Standards (Bio-Rad) were used as molecular weight markers. To stain LPS, the gels were stained using the Pro-Q Emerald 300 LPS gel staining kit according to the manufacturer’s instructions (Thermo Fischer Scientific). The presence of glycoproteins was examined by staining the gels with SYPRO Ruby protein gel stain according to the manufacturer’s instructions (Thermo Fisher Scientific). Stained LPS and protein bands were visualized under UV light using an Azure c300 gel imaging system (Azure Biosystems).

### Western blot analysis

Wild-type MVs (2 μg), Δ*spa* MVs (2 μg), and LPS (250 μg/mL) were either incubated with proteinase K (2 μg/mL) or with HEPES-NaCl buffer (at the same volume as proteinase K) for 30 min at 37°C. The protease reaction was stopped using 100 μg/mL phenylmethylsulfonyl fluoride (PMSF). The samples were then prepared and run on 4-20% SDS-PAGE gradient gels as described above. Precision Plus Protein Dual Color Standards (Bio-Rad) were used as molecular weight markers. After SDS-PAGE, samples on the gel were transferred onto a nitrocellulose membrane, followed by overnight incubation in blocking buffer at 4°C (Odyssey Intercept TBS Protein-free blocking buffer; Li-Cor). The membrane was then incubated for 1 h at RT with goat anti-Lipid A LPS antibodies (diluted 1:1,000 in blocking buffer; Bio-Rad; Cat # OBT1844). Blots were washed three times with TBST (50 mM Tris-Cl, pH 7.6, 150 mM NaCl, 0.1% Tween 20) followed by incubation at RT for 1 h with IRDye 680RD donkey anti-goat IgG secondary antibody (diluted 1:25,000 in blocking buffer; Li-Cor; Cat # 926-68074). After washing the membrane three times, the membranes were imaged on a Li-Cor Odyssey CLx near IR Imaging system (Li-Cor) with Li-Cor Image Studio software.

### Nuclease protection assays

To distinguish MV-associated nucleic acid molecules from extracellular nucleic acids not associated to MVs we conducted a series of nuclease protection assays. Five different conditions were prepared as follows: (i) 10 μL MVs (7 μg protein) were adjusted to 14 μL total volume with 4 μL HEPES-NaCl; (ii) 10 μL MVs were heated to 95 °C for 10 min, allowed to cool to RT, treated with 1 μL RIPA buffer (25 mM Tris-HCl pH 7.6, 150 mM NaCl, 1% NP-40, 1% sodium deoxycholate, 0.1% SDS) on ice for 10 min, and adjusted to 14 μL total volume with 3 μL HEPES-NaCl; (iii) 10 μL MVs were treated with 2 μL benzonase (250 U/μL) for 30 min at 37°C and the total volume was adjusted to 14 μL with 2 μL HEPES-NaCl; (iv) 10 μL MVs were heated, permeabilized with RIPA (1% final), treated with benzonase as described above and to total volume was adjusted to 14 μL with 1 μL HEPES-NaCl; (v) 10 μL MVs were heated, permeabilized with RIPA, treated with 1 μL Proteinase K (100 μg/mL final; Promega) incubated at 37°C for 30 min, the Proteinase K was inactivated with 1 μL PMSF (100 mM; Sigma-Aldrich) at RT for 5 min, and treated with 2 μL benzonase at 37°C for 30 min. These samples were degraded in 7 μL urea loading sample buffer, boiled for 5 min, and placed on ice for 2 min before loading on a 10% denaturing urea polyacrylamide gel in TBE buffer (Bio-Rad) next to an RNA marker (low range ssRNA ladder; NEB) and run at 200 V for 45 minutes. Staining with SYBR Gold (Invitrogen) allowed visualization of high and low molecular-weight MV-associated nucleic acids. To detect nuclease-treated MVs in native agarose gels, MV samples were mixed with 4 μL glycerol (20%) before being separated on a 2% TAE agarose gel, run at 110 V for 20 min, and stained with ethidium bromide (0.5 μg/mL).

### MV RNA & DNA extraction

Nucleic acids were extracted from 50 μL MV samples (28 μg total protein) using TRIzol. MVs were treated with HEPES-NaCl, benzonase, or Triton X-100 + Proteinase K + PMSF + benzonase as described above. After treatment, MVs were pelleted at 200,000 x g for 1h. MV pellets were then resuspended in 500 μL Trizol reagent and incubated for 5 min at room temperature. Next, 100 μL chloroform was added, mixed by inverting 3X and incubated for 3 min at RT. MVs were then centrifuged at 12,000 x g for 15 min in 4°C. The aqueous phase was transferred to a microcentrifuge tube (“MV RNA”) and the interphase layer was transferred to a microcentrifuge tube (“MV DNA”). MV DNA was kept on ice during the RNA extraction process. Each tube of MV RNA received 1 μL glycogen (20 mg/mL) and 250 μL isopropanol and were incubated for 10 min at 20°C. MV RNA tubes were centrifuged for 10 min at 12,000 x g at 4°C and the RNA pellet was washed with 500 μL ethanol (75%). The RNA pellet was resuspended in 20 μL nuclease-free dH_2_O and further ethanol precipitated. DNA was extracted from the MV DNA samples according to the TRIzol DNA Extraction protocol (Invitrogen). DNA pellets were resuspended in 8 mM NaOH adjusted to pH 7.0 with HEPES. RNA and DNA extracted from MVs were stored in 20°C. The quantity and size distribution of the RNA and DNA were determined by Agilent Bioanalyzer 2100 with an RNA 6000 Pico Kit and a DNA 1000 kit (Agilent Technologies, Santa Clara, CA, USA).

### Macrophage response assays

Macrophage cells were seeded into 12-well flat-bottom tissue culture plates at a volume of 1 mL per well (2 × 10^5^ cells/well) and incubated for 24 h at 37° C, 5% CO_2_. The following day, growth media was changed to fresh DMEM with 10% FBS and macrophages were treated with purified MVs at the indicated total protein concentration (1 μg/mL, 2 μg/mL, 5 μg/mL, 10 μg/mL or 20 μg/mL) or with HEPES-NaCl buffer (no treatment controls; NT) for 3 h. For time-course experiments, RAW 264.7 cells were treated with MVs (5 μg/mL) for the indicated time points. Following treatment with MVs or NT controls, the growth medium was removed from macrophages and the cells were homogenized with 500 μL TRIzol reagent (Invitrogen). RNA isolation was performed according to the manufacturer’s instructions.

For some experiments, RAW 264.7 cells were treated with 80 μM Dynasore hydrate (Sigma-Aldrich), 0.5 μM bafilomycin A1 (Sigma-Aldrich), 100 μM chloroquine (Sigma-Aldrich), 10 μM TLR3/dsRNA Complex Inhibitor (Millipore Sigma), or 0.1% DMSO for 30 minutes at 37°C, 5% CO_2_. Following pre-treatment with the aforementioned compounds, macrophages received the indicated MV dose or 10 μg/mL Polyinosinic-polycytidylic acid sodium salt [Poly (I:C)]. After 3 h, total RNA was extracted and processed for β-actin and IFN-β mRNA quantitation.

In control experiments, WT and *MyD88^-/-^/TRIF^-/-^* BMDMs were treated for 3 h at 37°C, 5% CO_2_ with 1 μg/mL 2’3-cGAMP (negative control; Sigma-Aldrich) or with 2 μM CpG ODN 2395 (positive control) 5’-T*C*G*T*C*G*T*T*T*T*C*G*G*C*G*C*G*C*G*C*C*G-3’ (*represents phosphorothioate-modified bases; Eurofins). To confirm that TLR3, TLR7, TLR9 and IRF3/7-deficient macrophages produce significantly less IFN-β mRNA in response to known TLR ligands, we performed a series of control experiments. WT and *TLR3^-/-^* macrophages were treated with 20 μg/mL Poly (I:C) alone or Poly (I:C) was transfected into cells by using Lipofectamine 2000 (Invitrogen) as the transfection reagent. Transfection was conducted as follows: 40 μL Poly (I:C) (10 mg/mL) was diluted in 160 μL Opti-MEM reduced serum medium (Thermo Fisher) and mixed gently to obtain a final concentration of 2 mg/mL Poly (I:C). 4 μL Lipofectamine 2000 was mixed with 196 μL Opti-MEM and incubated at RT for 5 min. The diluted Poly (I:C) and Lipofectamine 2000 samples were combined, mixed gently, and incubated for 20 min at RT. Mock transfection reagent was prepared as described above, except Poly (I:C) was replaced with dH_2_O. 20 μL Poly (I:C)+Lipofectamine [1 mg/mL Poly (I:C)] or 20 μL mock transfection reagent was added to WT and *TLR3^-/-^* cells to a final concentration of 20 μg/mL Poly (I:C). WT and *TLR^-/-^* cells were treated with 1 μg/mL Resiquimod, WT and *TLR9^-/-^* cells were treated with 5 μM CpG ODN 2395 for 3h or were transfected with 5 μM CpG ODN 2395 complexed with 1 μL Lipofectamine 2000 (in 100 μL Opti-MEM) for 3h, and WT and *IRF3^-/-^ IRR^-/-^* macrophages were treated with 20 μg/mL Poly (I:C)+Lipofectamine for 3 h. Total RNA was extracted from these cells and processed for qPCR analysis.

For *S. aureus* RNA transfection assays, 60 μL of total RNA (1 μg) was diluted in 50 μL Opti-MEM. 1 μL Lipofectamine 2000 was mixed with 49 μL Opti-MEM. The mock transfection reagent control was prepared by replacing RNA with dH_2_O. Transfection was carried out in RAW 264.7 cells according to the manufacturer’s instructions. After 3h, macrophages were harvested, and the cellular RNA was extracted and analyzed for β-actin and IFN-β mRNA quantitation.

### Macrophage response assays with nuclease treated MVs

To assess the response of RAW264.7 macrophages to stimulation with intact MV compared to stimulation with nuclease-treated MVs, MVs were treated with benzonase (Sigma-Aldrich) at a ratio of 10 μg MV total protein:1 μL benzonase (250 U/μL) for 30 min at 37°C. To control for functional effects that may be caused by this incubation period, intact MVs were treated with HEPES-NaCl at the same ratio of benzonase volume to total MV protein. We also prepared a HEPES-NaCl + 1 μL benzonase sample for NT controls and both intact MVs and NT control samples were incubated for 30 min at 37°C. To remove benzonase from MVs prior to macrophage stimulation assays, benzonase-treated MVs, intact MVs, and NT control samples were ultracentrifuged at 200,000 × g for 1h. MV pellets were then resuspended in HEPES-NaCl and were added to RAW264.7 cells at the indicated doses for 3 h.

### cDNA synthesis and qPCR

cDNA was synthesized using the High-Capacity cDNA Reverse Transcription Kit (Applied Biosystems) in accordance with the manufacturer’s instructions. The reaction mixture consisted of 500 ng total RNA, 2 μL RT buffer, 2 μL Reverse Transcriptase (RT) random primers, 0.8 μL dNTP Mix (100 mM), and 1 μL of MultiScribe Reverse Transcriptase (50 U/μL) in a final reaction volume of 20 μL. The synthesized cDNA was diluted 20X to 25 ng in RNase-free dH_2_O.

The reaction mixture for quantitative real-time PCR (qPCR) consisted of 1 μL template cDNA (25 ng), 10 μL PowerUp SYBR Green Master Mix (Thermo Fisher Scientific), and 3 μL of each primer pair (300 nM final primer concentration) in a total reaction volume of 20 μL. qPCR was performed on a StepOnePlus Real-Time PCR System (Applied Biosystems) to detect mouse IFN-*α*2, IFN-*α*4, IFN-β, and β-actin The qPCR primers used in this study are experimentally validated sequences taken from the Primer Bank database (https://pga.mgh.harvard.edu/primerbank/). The following primer pairs were amplified using qPCR: IFN-*α* 2 gene forward primer, 5’-TACTCAGCAGACCTTGAACCT-3’; reverse primer, 5’-CAGTCTTGGCAGCAAGTTGAC-3’; IFN-*α*4 gene forward primer, 5’-TGATGAGCTACTACTGGTCAG C-3’; reverse primer, 5’-GATCTCTTAGCACAAGGATGGC-3’; IFN-*β* gene forward primer, 5’-AGCTCCAAGAAAGGACGAACA-3’; reverse primer, 5‘-GCCCTGTAGGTGAGGTTGAT-3’; β-actin gene forward primer, 5’ GGCTGTATTCCCCTCCATCG-3’; reverse primer, 5’ CCAGTTGGTAAC AATGCCATGT-3’.

The mRNA expression levels for IFN-*α*2, IFN-*α*4, or IFN-*β* with respect to β-actin were calculated using the delta-delta cycle threshold (Ct) method (2^-ΔΔCt^) and the standard error of the difference between means were determined using Microsoft Excel.

### Microscopy imaging of RAW 264.7 cells

From confluent RAW 264.7 cells, 2×10^5^ macrophage cells were plated in 35mm dishes and incubated at 37°C, 5% CO_2_ for 24 h. The next day, MVs (1 μg/mL) were labeled with 1 μL SYTO RNASelect (1 μM). Unincorporated dye was removed by washing MVs in HEPES-NaCl 2X after ultracentrifugation at 200,000 for 1 h. The RAW 264.7 growth media was changed to fresh DMEM with 10% FBS and macrophages were treated with purified MVs (10 μg/mL protein). After 10 minutes of stimulation, macrophages were washed with 1X PBS and fixed with 4% paraformaldehyde. For immunostaining, cells were permeabilized with warmed 0.1% Triton X-100 in PBS for 5 min (this step was repeated 3X). Cells were then incubated in blocking solution (1% bovine serum albumin in PBS) overnight at 4°C. The following day, the cells were washed in PBS and incubated for 1-h at RT with 1 mL anti-Early Endosomal Antigen 1 (EEA1) monoclonal antibody (F. 43.1; Invitrogen) diluted 1:200 in blocking buffer. Cells were then rinsed for 5 min at RT (3X) with PBS and incubated with secondary antibody (Alexa Fluor 647 diluted 1:2,000 in blocking buffer; Invitrogen) for 45 min at RT. The cells were then rinsed 3X for 5 min in 1X PBS and imaged using an N–STORM super resolution microscope (Nikon). The extent of colocalization between MV-RNA and the endosomal marker EEA1 was calculated using the CoLocalizer Pro 5.2 software with the Mander’s overlap coefficient algorithm (Colocalization Research Software, Switzerland, Japan).

### 5-ethynyl uracil (EU) labeling and imaging of bacterial RNA

To label *S. aureus* cells with 5-EU *in vivo,* we grew a 5 mL overnight bacterial culture at 37°C with shaking (200 rpm) in RNased NB. The following day, the cultured cells were diluted 1:500 by adding 20 μL of the cell suspension to a flask containing 10 mL RNased NB and 10 μL 5-EU (100 mM stock solution; Invitrogen), giving a final concentration of 100 μM 5-EU. Cells were grown to OD_600_ of 1 (~5 h) and 1 mL of the cultured cells was removed and centrifuged at 18,000 x g for 5 min. Cell pellets were washed in PBS and fixed using 500 μL of fixation solution (4% paraformaldehyde in PBS; Sigma-Aldrich) for 15 min at RT followed by a 15 min incubation on ice. Fixed cells were washed 3X in PBS and permeabilized by resuspending the fixed cell pellet in 200 μL lysostaphin (80 μg/mL solution in TE buffer) and incubating at 37°C for 15 min. 100 μL of permeabilized cells were pipetted onto coverslips coated with 0.01% poly-L-lysine (Sigma-Aldrich). Cells were allowed to adhere to the coverslip at RT for 15 min and unbound cells were washed with PBS. To detect 5-EU labeled RNA by fluorescence microscopy, the commercially available kit, Click-iT RNA Alexa Fluor 488 Imaging Kit (Invitrogen) was used. The Click-iT reaction cocktail was prepared and added onto coverslip-adhered cells according to the manufacturer’s instructions. After removing the Click-iT reaction rinse buffer, coverslips were washed once more with PBS (protected from light). The coverslips were then mounted onto a clean microscope slide using ProLong Diamond Antifade Mountant with DAPI (Invitrogen). CSLM was carried out using a Zeiss LSM 880 inverted confocal microscope and images were acquired using a 40x water immersion objective. Image acquisition of 5-EU Alexa Fluor 488-labeled cells was carried out with the 488 nm Argon laser, the 405 nm Diode laser was used to acquire signals from DAPI-stained cells, and brightfield images were captured using the transmitted light detector (T-PMT). Acquired images were processed and analyzed using the ZEN Blue 2.3 software (Zeiss).

### 5-EU labeling of MV-associated RNA and DiD-MV labeling

5-EU labeling of *S. aureus* RNA for MV isolation was performed as described above, except cells were grown in a flask containing 250 mL RNased NB supplemented with 150 μM 5-EU. After reaching mid-log phase (OD_600_ of 1.0), MVs were isolated from the supernatant of the culture as delineated in the *“Membrane Vesicle Isolation”* section. However, after the initial filtration step, supernatants were pelleted using a Beckman Avanti J-25 centrifuge (38,400 x g, 3h, 4°C; JLA-16.250 rotor), followed by ultracentrifugation and filtration using a 0.22-μm pore size, hydrophilic polyvinylidene fluoride filter (Merck Millipore Germany). MV preparations were tested for sterility and the protein concentration was measured using a Bradford assay.

For MV RNA uptake assays, 5-EU RNA MVs were treated with benzonase as described in the *“Macrophage response assays with nuclease-treated MVs”* section. Following benzonase treatment, bMVs were labeled with 5% (v/v) DiD Vybrant cell-labeling solution (Molecular Probes). Specifically, 47.5 μL bMVs were incubated with 2.5 μL DiD (5 μM) for 30 min at 37°C (protected from light). Unbound dye was removed by ultracentrifugation (200,000 x g, 30 min, 4°C), MVs were washed in PBS, and subjected to ultracentrifugation once more. DiD-labeled 5-EU RNA bMV pellets were resuspended in 47.5 μL PBS and stored at −20 °C.

bMVs purified from *S. aureus* cells grown in the absence of 5-EU were also stained with 5% (v/v) DiD and with 5% (v/v) SYTO RNASelect working stock solution (5 μM). bMVs were then incubated at 37°C for 30 min, and unincorporated dye was removed by ultracentrifugation 2X. DiD + SYTO RNA-labeled bMVs were resuspended in PBS and stored at −20 °C.

### Airyscan super-resolution microscopy imaging of MV uptake by macrophages

RAW 264.7 cells or wild-type macrophages (2 x 10^5^ cells/well in 6-well plates) were grown on poly-L-lysine-coated glass coverslips in DMEM supplemented with 10% FBS. After 24 h, macrophages were treated with 5 μg/mL unlabeled bMVs, SYTO RNA-labeled bMVs, DiD-labeled 5-EU RNA bMVs, or with DiD + SYTO RNA-labeled bMVs for 1 h. Macrophages were then washed with PBS, fixed with pre-warmed 4 % paraformaldehyde in 1X PBS for 15 min at RT. Fixed cells were washed with PBS 3X and permeabilized with 0.2% saponin (Sigma-Aldrich) for 10 min at RT. After washing 3 times with PBS, 5-EU RNA bMV-treated macrophages were incubated with the Click-iT reaction cocktail containing CuSO4 and Alexa Fluor 488 azide to label internalized bMV-5-EU RNA according to the manufacturer’s instructions. For colocalization assays of bMV-RNA in RAW 264.7 cells, the permeabilized macrophages were incubated in blocking buffer (1% BSA in PBS) with or without human monoclonal IgG1 kappa isotype control antibody (10 μg/mL; GeneTex; Cat # GTX35068) for 2 h at RT. The macrophages were then incubated with EEA1 antibodies conjugated to Alexa Fluor 647 (1:100 dilution in blocking buffer; Santa Cruz Biotechnology; Cat # sc-137130) for 1 h at RT. The coverslips were washed three times in PBS and mounted onto slides in ProLong Diamond Antifade Mountant with DAPI.

All samples were imaged using a Zeiss LSM 880 inverted confocal equipped with an Airyscan detection unit (63x/1.4 oil immersion objective). DAPI was excited at 405 nm, Alexa Fluor 488 and SYTO RNASelect were excited at 488 nm, DiD or EEA1-Alexa Fluor 647 were excited at 633 nm, and brightfield images were captured using the T-PMT. After acquisition of selected cells using T-PMT in confocal mode, cells were imaged in super-resolution mode with the Airyscan detector (pinhole size 1.25 AU). The z-stack module was used to capture between 40 and 64 sections per z-stack (6.209 μm and 10.045 μm, respectively) at 0.16 μm intervals per slice. These intervals were automatically calculated by the Zen Black software to fulfill Nyquist criteria. Consecutive Z-stack images captured by Airyscan were reconstructed to a 3D image using the Zen Blue 2.3 software (using maximum intensity projection).

## Acknowledgements

We thank Dr. Jörn Coers for kindly providing the bone-marrow derived macrophages, Dr. Ken Yokoyama for providing the *S. aureus* Newman strain, and Dr. Terry Oas for providing the *S. aureus* Newman Δ*spa* strain. The following reagents were obtained through BEI Resources, NIAID, NIH: Macrophage Cell Lines Derived from Wild Type Mice, NR-9456; IRF3/IRF7 Double Knockout Mice, NR-15637; TLR3 Knockout Mice, NR-19974; TLR7 Knockout Mice, NR-15634; and TLR9 Knockout Mice, NR-9569. We thank Clariss Limso for assistance with the LIVE/DEAD assays. We also acknowledge the support and the use of resources of the Duke Shared Materials Instrumentation Facility for TEM imaging and the use and the Duke Light Microscopy Core Facility for use of the confocal and super-resolution microscopes. This research was supported by Duke University Medical Center, the Burroughs Wellcome Fund Graduate Diversity Enrichment Program, Duke Cancer Research Education Program Scholar Awards, and NSF IOS-1931309.

## Author contributions statement

B.V.R. and M.J.K. conceived the experiments, B.V.R. conducted the experiments, B.V.R. and M.J.K. analyzed data, wrote, and reviewed the manuscript.

## Additional information

The authors have no competing interests.

