## Supplemental Figures for "*Staphylococcus aureus* secretes immunomodulatory RNA and DNA via membrane vesicles"

**Supplementary File**

**(Figures S1-S18)**

**for**

### "*Staphylococcus aureus* secretes immunomodulatory RNA and DNA via membrane vesicles"


**Blanca V. Rodriguez and Meta J. Kuehn***

Duke University, Department of Biochemistry, Durham, 27710, USA
*

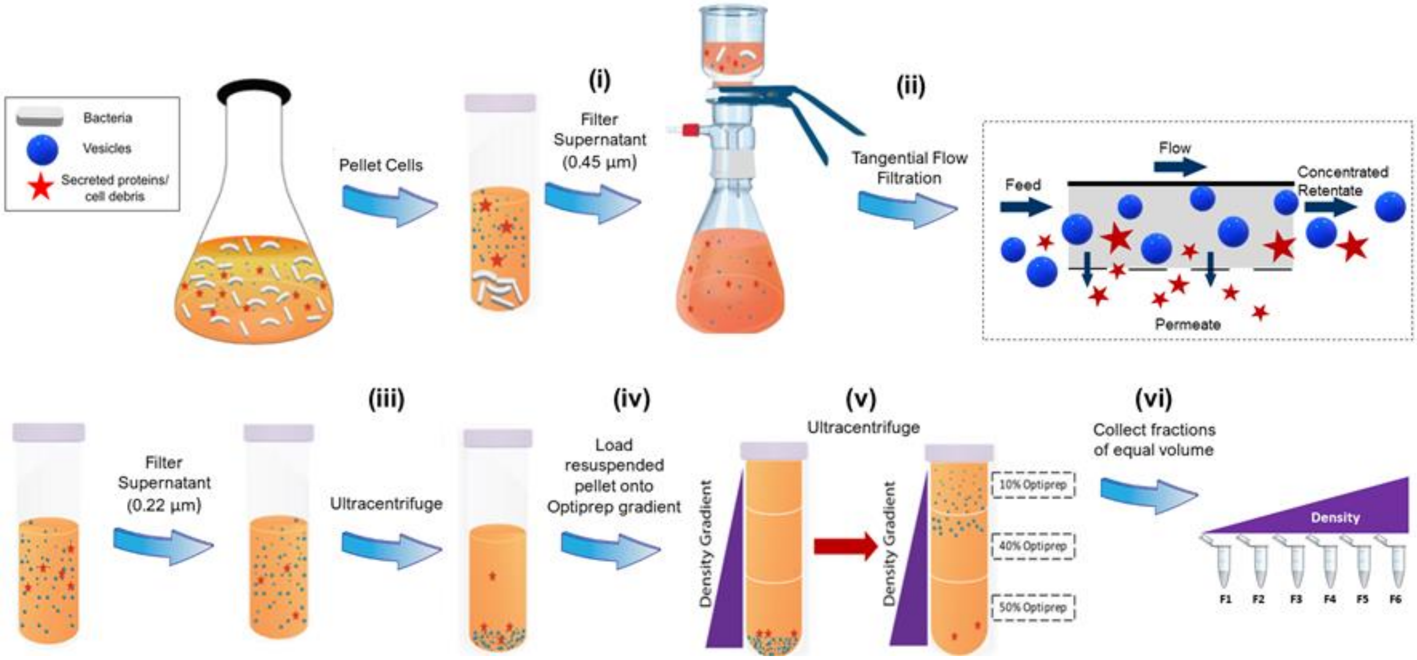


**Figure S1: Schematic representation of MV isolation workflow.** *S. aureus* was cultivated in nutrient broth to an $OD_{600}$ of 1.0 and the intact bacteria was sedimented with gentle centrifugation. (i) The supernatant was collected and filtered to remove cells. (ii) The supernatant was concentrated (12X) using a Tangential Flow Filtration system with a 100 kDa-cutoff filter and the retentate was subsequently filtered through a 0.22 µm pore. (iii) The concentrated filtrate was pelleted at high speed (iv) and the vesicle pellet resuspended in Optiprep. (v) The resuspended pellet was placed in the bottom of a 12.5 mL ultracentrifuge tube and overlaid with sequential decreasing percentages of Optiprep solutions. (vi) After ultracentrifugation, fractions were collected starting from the top of the gradient (F1-F6).

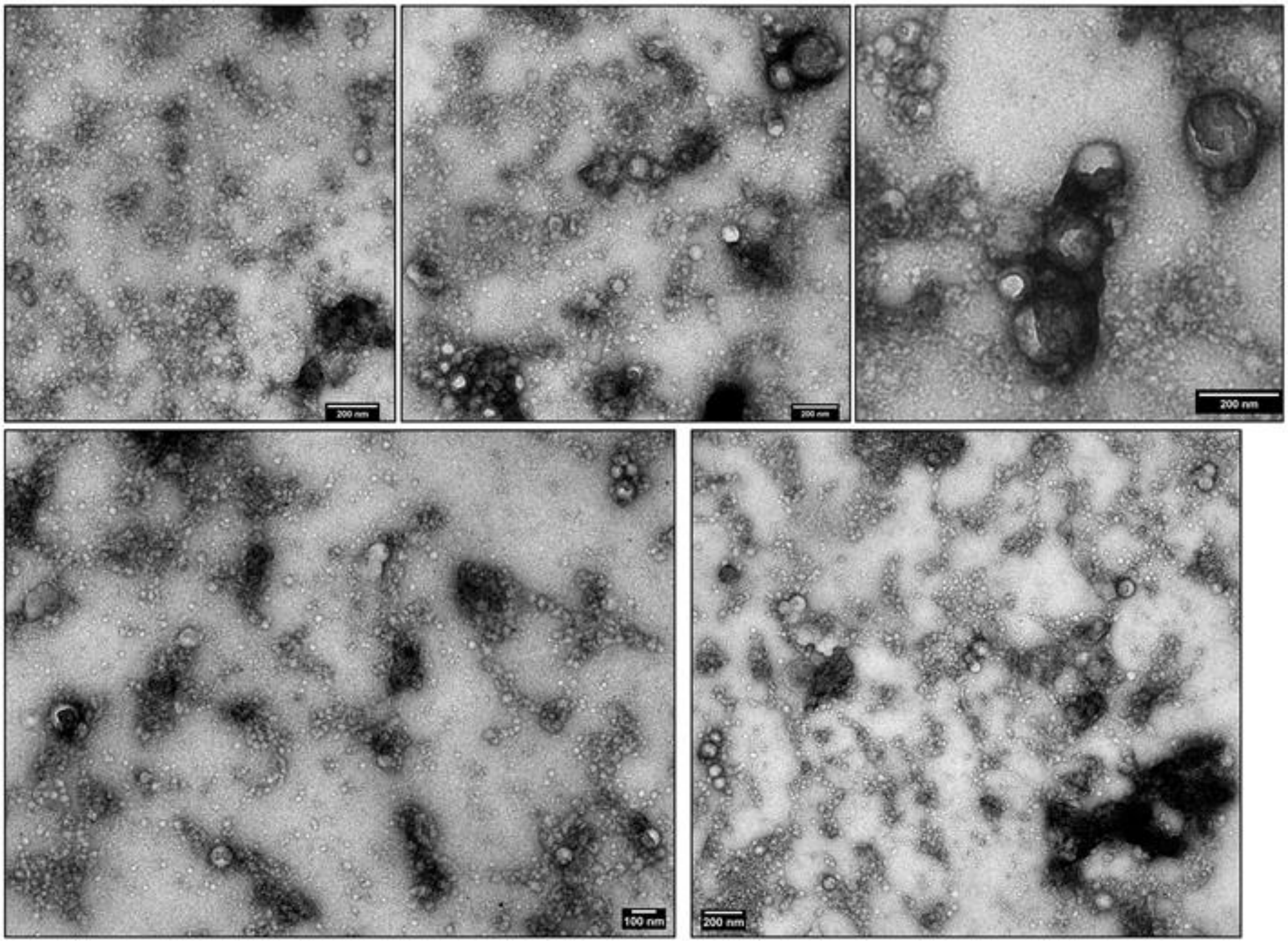


**Figure S2: TEM images of purified MVs.** Representative micrographs of negatively stained MVs collected from *S. aureus* Newman after Optiprep density gradient purification. Scale bars are all 200 nm except for in the bottom left panel where it indicates 100 nm.

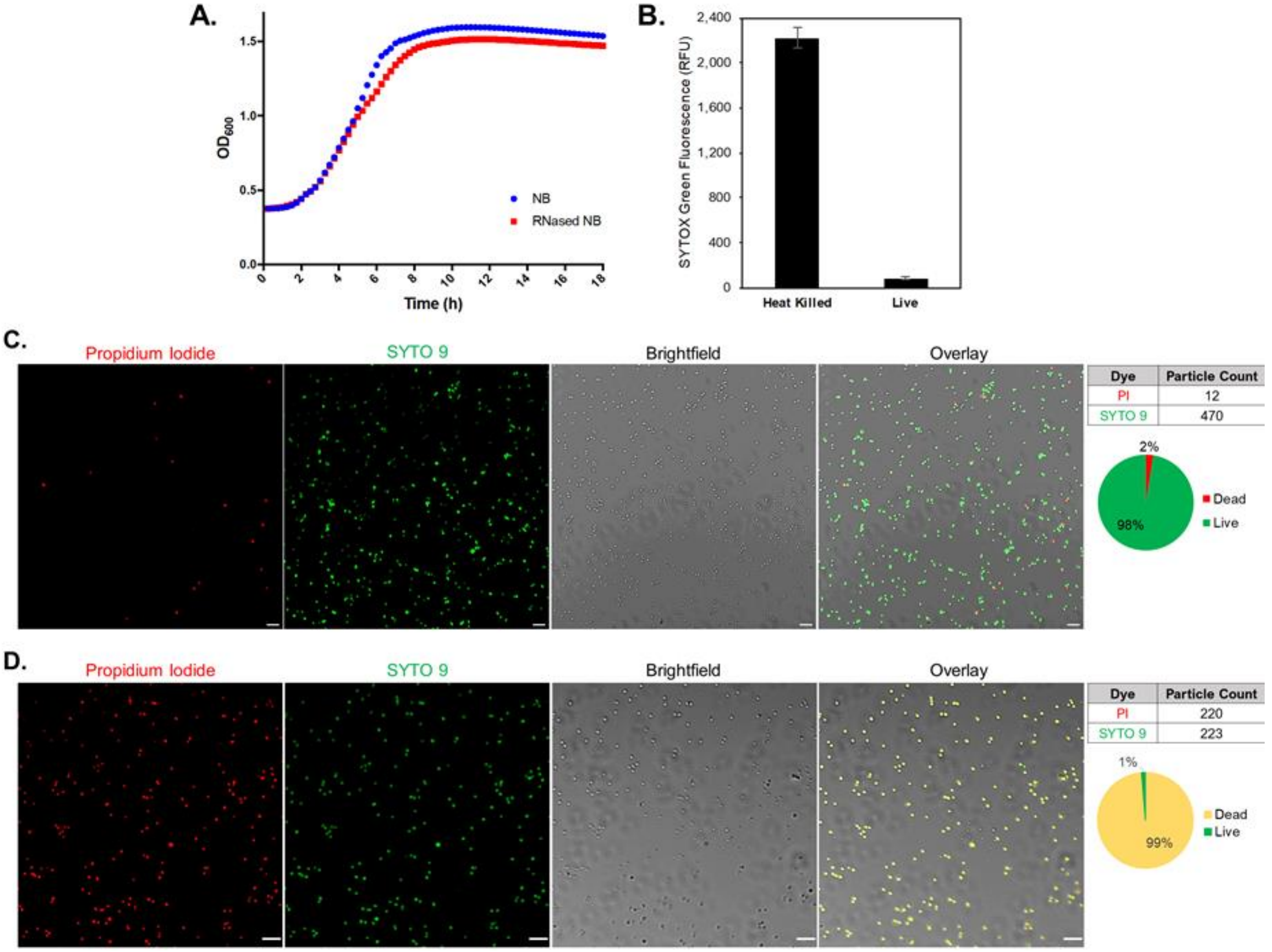


**Figure S3: Minimal cell lysis occurs at *S. aureus* mid-log growth phase.** (A) Representative growth curve of *S. aureus* Newman grown in nutrient broth (NB) treated with or without RNase A over the course of 18 hours. Bacterial cultures were grown overnight in NB at 37˚C with aeration. After 18h, cultures were inoculated in NB treated with or without RNase A (0.5 µg/mL) in a 1:100 dilution. Optical densities were recorded during the subsequent incubation (shaking, 37˚C) over an 18 h period. Representative growth curves from triplicate experiments are shown. (B) SYTOX Green relative fluorescence units (RFU) of heat-killed or mid-log phase *S. aureus* cell normalized to mean negative controls (PBS). (C-D) Confocal microscopy images of live (C) and (D) isopropanol-killed *S. aureus* Newman cells grown to early-stationary phase ($OD_{600}$ 1.6). Cells were prepared using a LIVE/DEAD BacLight Bacterial Viability Kit followed by staining with SYTO9 (green fluorescence for live bacteria) and propidium iodide, PI, (red fluorescence for killed bacteria) for 15 min in the dark at 25°C. PI and SYTO9-stained cell counts were obtained using the Analyze Particles function available in ImageJ. Scale bars = 5 µm.

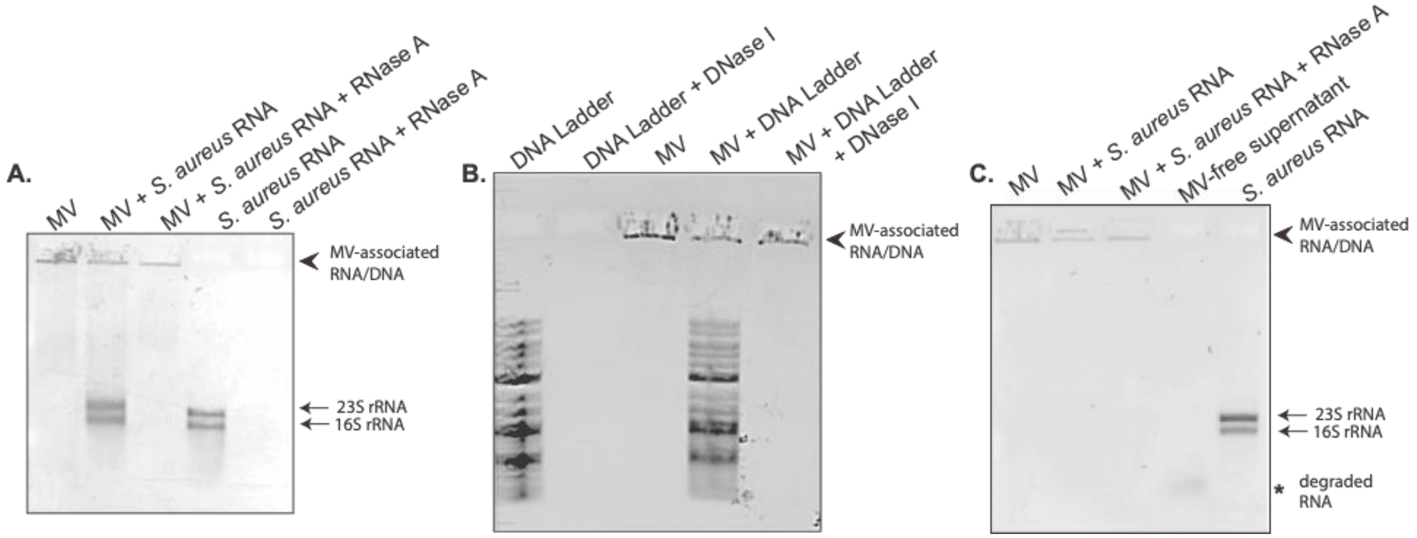


**Figure S4: MVs do not readily bind exogenous RNA or DNA.** (A) RNA extracted from whole-cell *S. aureus* was incubated with native MVs for 30 min at RT, followed by RNase A treatment at 37˚C for 30 min as indicated. MV + RNA mixtures were loaded directly onto a 1% TAE agarose gel and stained with ethidium bromide. Samples of MVs, *S. aureus* RNA, and *S. aureus* RNA incubated with RNase A were loaded as controls. (B) A sample of 2-log DNA ladder was incubated with native MVs for 30 min at RT, followed by digestion with DNase I for 30 min at 37˚C as indicated. MV + DNA ladder mixtures were loaded directly onto a 1% TAE agarose gel and stained with ethidium bromide. Samples of DNA ladder, DNA ladder incubated with DNase, and MVs were loaded as controls. (C) Native MVs + *S. aureus* RNA were mixed and incubated at RT for 30 min, centrifuged for 1 h at 200,000 x g to remove unbound exogenous RNA from MVs, pellets were resuspended in HEPES-NaCl. Samples were then loaded onto 1% TAE agarose gels and stained with ethidium bromide. Arrowhead indicates migration of nucleic acid associated with intact MVs. Asterisk indicates unbound and degraded RNA that had been exogenously added to the MVs. Arrows indicate the migration of 23S and 16S rRNA from whole cell *S. aureus* RNA preparations.

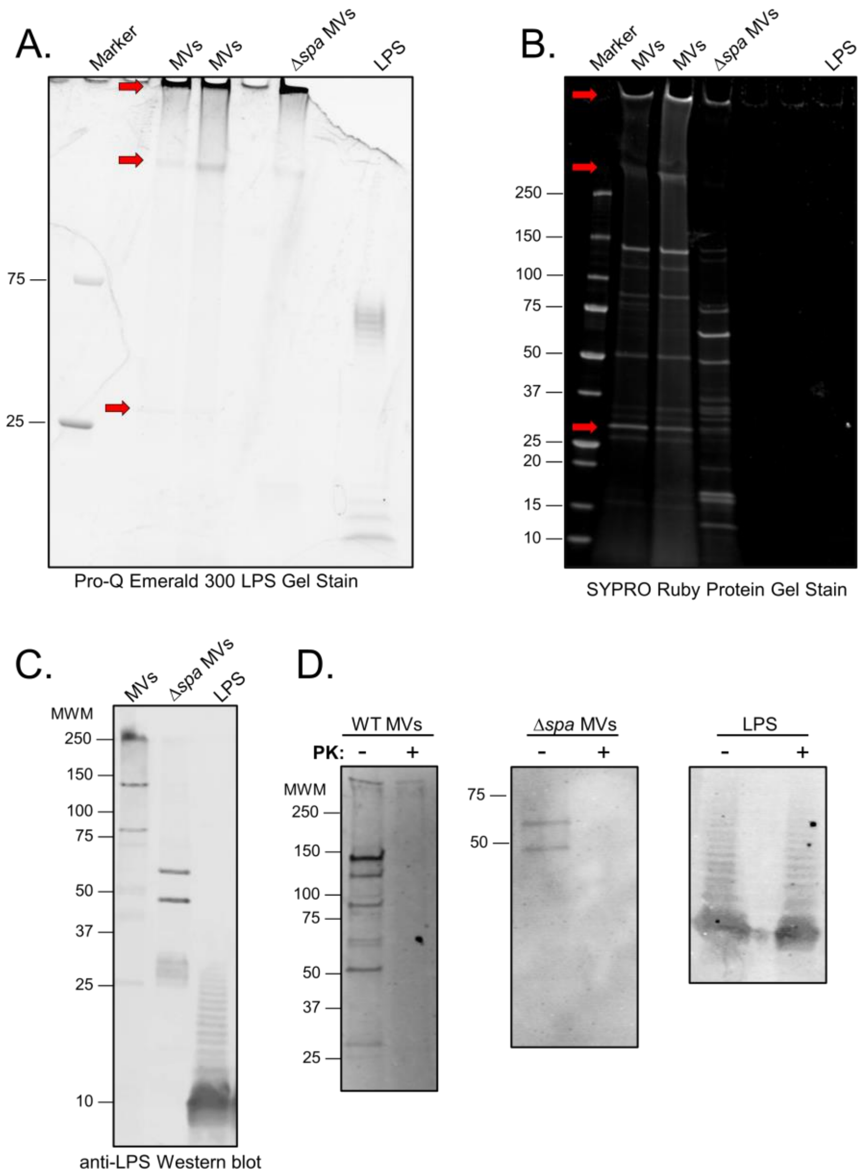


**Figure S5: *S. aureus* MV samples are free of endotoxin contamination.** Wild-type MVs (samples from two different vesicle purifications are shown), *Δspa* MVs, and LPS standard from *E. coli* serotype 055:B5 were separated on a 4-20% SDS-PAGE gradient gel. Gels were stained using the (A) Pro-Q Emerald 300 LPS Gel Stain or the (B) SYPRO Ruby Protein Gel Stain. Glycoproteins are indicated by red arrows. (C, D) After treatment of WT MVs, *Δspa* MVs, and LPS with or without Proteinase K (PK), proteins were separated by SDS-PAGE and transferred to a nitrocellulose membrane. LPS was detected in the samples by immunoblotting with polyclonal antibodies against the endotoxin region of LPS. Molecular weight marker (MWM, in kDa).

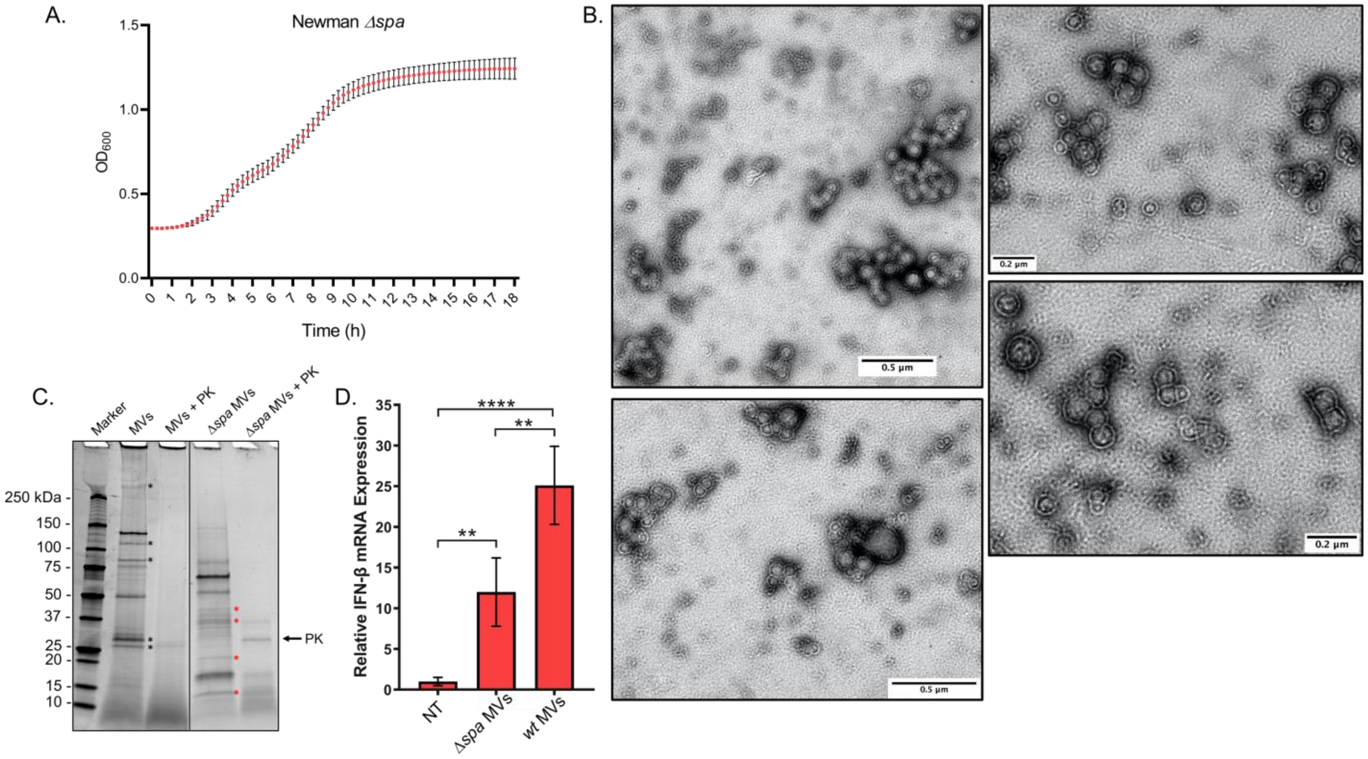


**Figure S6: Protein A may contribute to the immunomodulatory activity of MVs.** (A) Growth curve of Newman *Δspa* cells in RNased-NB media at 37°C. The data is represented as mean ±SE from n = 7. (B) MVs purified from Newman *Δspa* were imaged using negative-stain TEM. Scale bars are indicated in each micrograph. (C) The protein profile of MVs purified from wild-type Newman and *Δspa* were examined using SDS-PAGE. MVs were treated with or without Proteinase K (2 µg/mL) for 30 min at 37° before being separated on a 4-20% SDS-PAGE gradient gel. Protein gels were stained using SYPRO Ruby Protein Gel Stain. (D) Wild-type macrophage cells were treated with HEPES-NaCl (NT), *Δspa* MVs (5 µg/mL), or MVs (5 µg/mL). After 3 h, macrophages were homogenized for qPCR analysis of IFN-β mRNA expression relative to β-actin mRNA expression. The data is represented as mean ±SE from n = 3. Statistical analysis was performed by one-way ANOVA with Tukey's multiple comparison test (**$p < 0.005$; ****$p < 0.0001$).

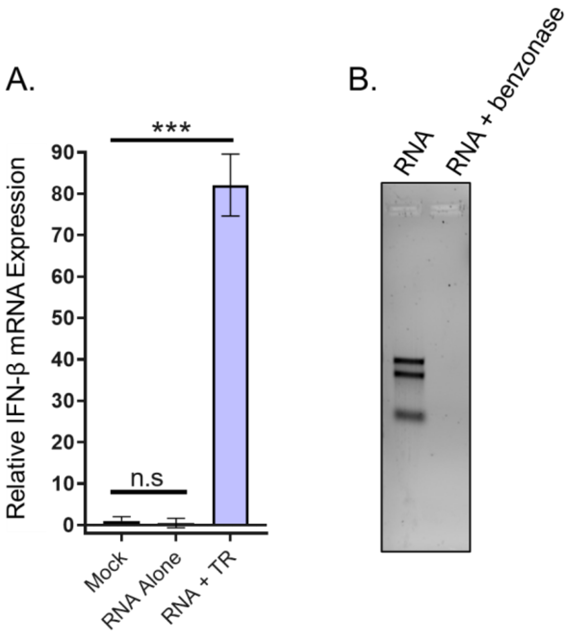


**Figure S7: *S. aureus* total RNA induces IFN-β mRNA expression in macrophages only after transfection and is benzonase sensitive.** (A) Total RNA (1 µg) extracted from *S. aureus* was coincubated with RAW 264.7 macrophages with or without Lipofectamine (‘RNA alone’ = without Lipofectamine; ‘RNA + TR’ = with Lipofectamine). Macrophages were also treated with transfection reagent alone (mock). After 3 h, macrophages were homogenized for qPCR analysis of IFN-β mRNA expression relative to β-actin mRNA expression. The data is represented as mean ±SE from n = 2. Statistical analysis was performed by one-way ANOVA with Tukey’s multiple comparison test (n.s = not significant; ***p < 0.001). (B) *S. aureus* RNA (400 ng) without or with benzonase treatment was run on a 0.5% agarose gel and stained with ethidium bromide.

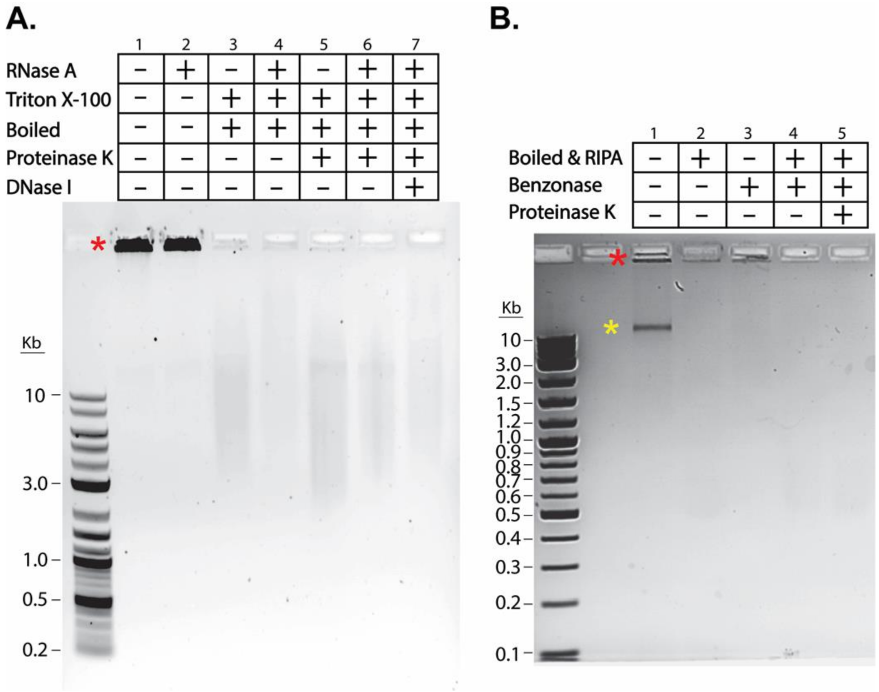


**Figure S8: Subpopulations of MV-associated nucleic acids are protected from RNase & DNase degradation irrespective of MV permeabilization and/or Proteinase K treatment.** (A) MV samples were treated for 30 min with either HEPES-NaCl [1], RNase A at 37˚C [2] or with the detergent Triton X-100 for 10 min at 4˚C followed by boiling for 10 min at 90˚C [3]. MV sample was treated with RNase A followed by detergent and heating [4], or detergent, heat, and Proteinase K [5], or similar to 5 with the addition of RNase A [6]. MV sample was treated with all of the aforementioned conditions [7]. (B) MVs were treated with HEPES-NaCl [1], with the detergent RIPA followed by boiling [2], or with Benzonase for 30 min at 37˚C [3]. MV sample was treated with detergent, heat, and Benzonase [4]. MV sample was treated with detergent, heat, Benzonase, and Proteinase K [5]. Subsequently, the MV samples were loaded onto 1% TAE agarose gels and stained with ethidium bromide. Red asterisks indicate nucleic acids that remain in the gel loading wells and the yellow asterisk indicates a band of Benzonase-sensitive nucleic acids.

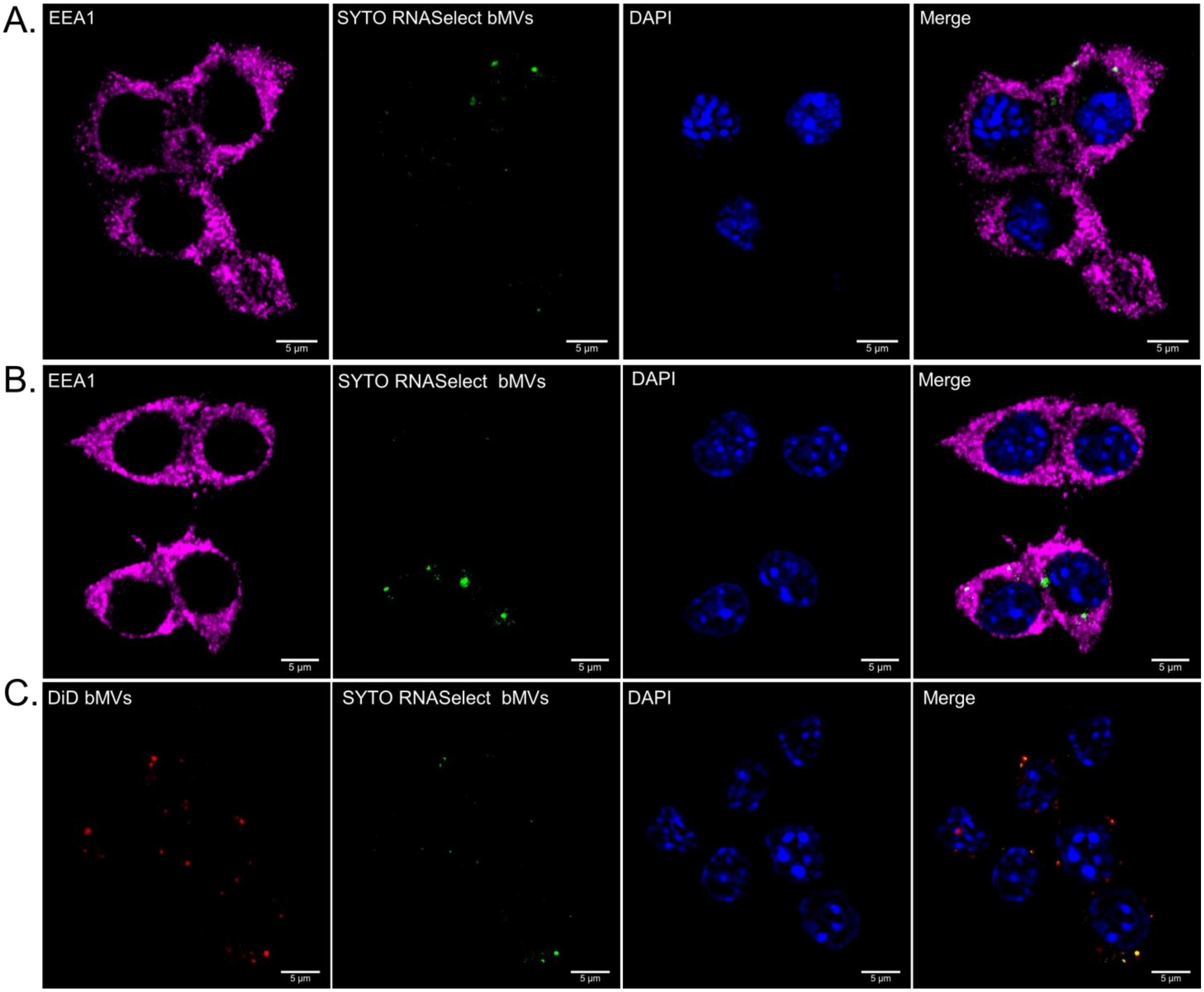


**Figure S9: Airyscan super-resolution microscopy shows moderate colocalization of bMV-associated RNA to early endosomal compartments in RAW 264.7 macrophages.** RAW 264.7 macrophages were treated with 1 µg/mL SYTO RNASelect-labeled bMVs (green) for 10 min. Macrophages were then fixed and permeabilized using saponin. Macrophages were either incubated with blocking buffer alone (A) or with blocking buffer containing an irrelevant human IgG monoclonal antibody (B). Cells were then immunostained with anti-EEA1 Alexa Fluor 647 antibodies (magenta), washed, and counterstained with DAPI (nucleus; blue). Representative z-stack projections are shown from 2 independent experiments. (C) Macrophages were treated with DiD + SYTO RNA-labeled bMVs (green; bMV RNA & magenta; bMV lipids) for 10 min. Macrophages were then fixed and stained with DAPI. Representative z-stack projections are shown from 2 independent experiments. Images demonstrate that bMV-associated RNA molecules can be delivered to early endosomes in macrophages after 10 min of coincubation. All images were captured using a Zeiss LSM 880 equipped with Airyscan (63x objective). Scale bars = 5 µm.

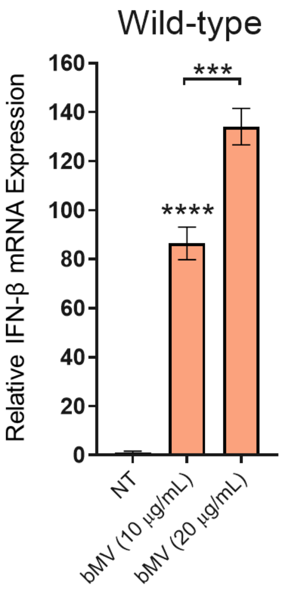


**Figure S10: Wild-type macrophages produce IFN-β after stimulation with bMVs in a dose-dependent manner.** WT macrophage cells were treated with HEPES-NaCl (NT), 10 µg/mL bMVs, or 20 µg/mL bMVs. After 3 h, macrophages were homogenized for qPCR analysis of IFN-β mRNA expression relative to β-actin mRNA expression. The data is represented as mean ±SE from $n = 3$. Statistical analysis was performed by one-way ANOVA with Tukey's multiple comparison test (***$p < 0.001$; ****$p < 0.0001$).

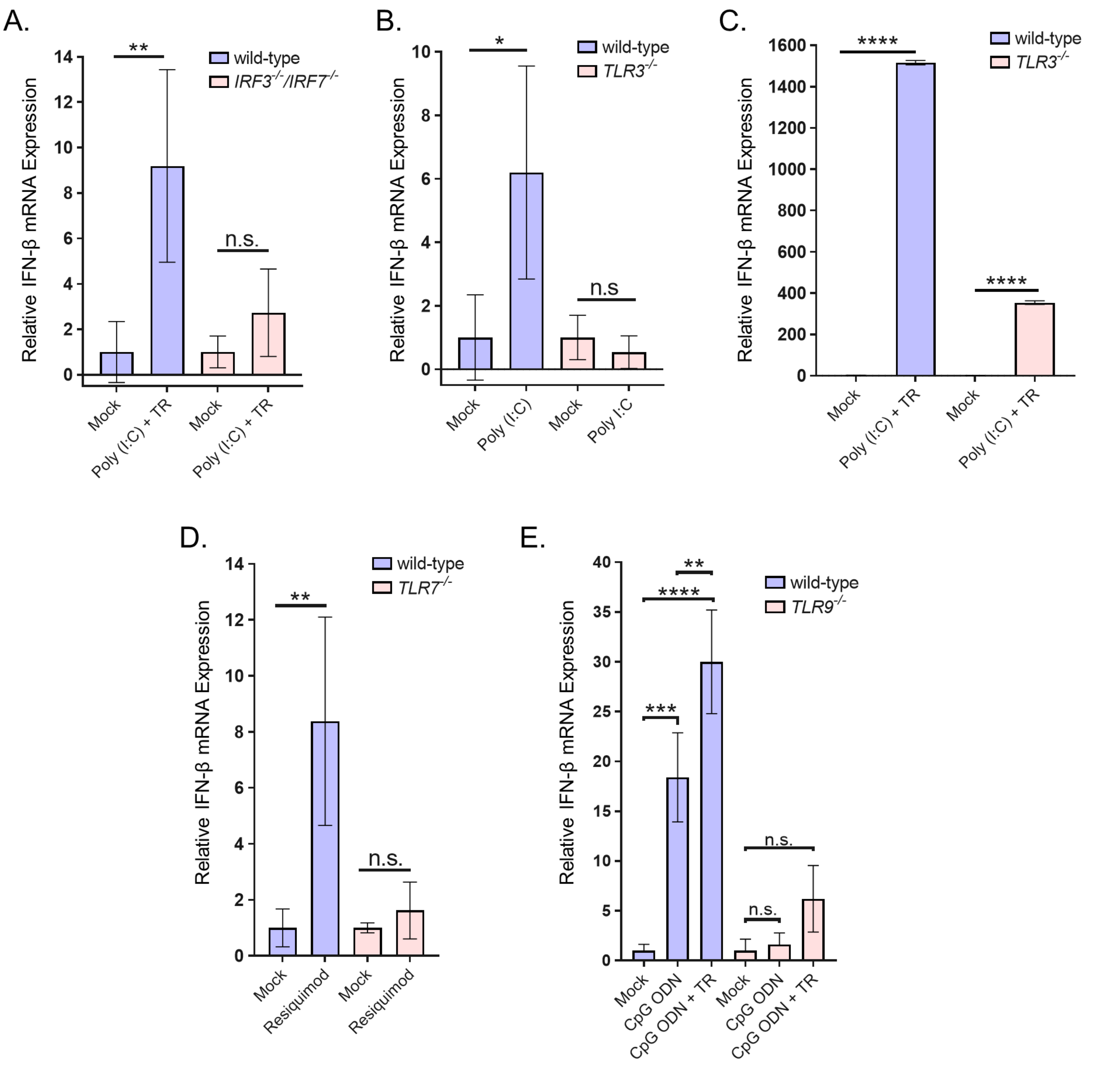


**Figure S11: Control experiments show that IRF3/7, TLR3, TLR7, and TLR9-deficient macrophages produce significantly less IFN-β in response to known TLR ligands relative to WT macrophages.** (A) WT and *IRF3*$^{-/-}$*IRF7*$^{-/-}$ macrophages were treated with mock transfection reagents or with 20 µg/mL Poly (I:C)+Lipofectamine mixtures for 3 h. (B) *TLR3*$^{-/-}$ and WT macrophages were treated with 20 µg/mL Poly (I:C) or mock reagents for 3 h. (C) *TLR3*$^{-/-}$ and WT macrophages were treated exactly as described for *IRF3*$^{-/-}$*IRF7*$^{-/-}$ macrophages. (D) WT and *TLR7*$^{-/-}$ macrophages were treated with 1 µg/mL Resiquimod for 3 h. (E) WT and *TLR9*$^{-/-}$ cells were treated with 5 µM CpG ODN 2395 or with 5 µM CpG ODN+Lipofectamine for 3 h. After stimulation with their corresponding ligands, macrophages were harvested for qPCR analysis of IFN-β mRNA expression relative to β-actin mRNA expression. The data is represented as mean ±SE, n = 3. Statistical analysis was performed by one-way ANOVA with Tukey's multiple comparison test (*$p \leq 0.05$; **$p < 0.005$; ***$p < 0.001$; ****$p < 0.0001$).

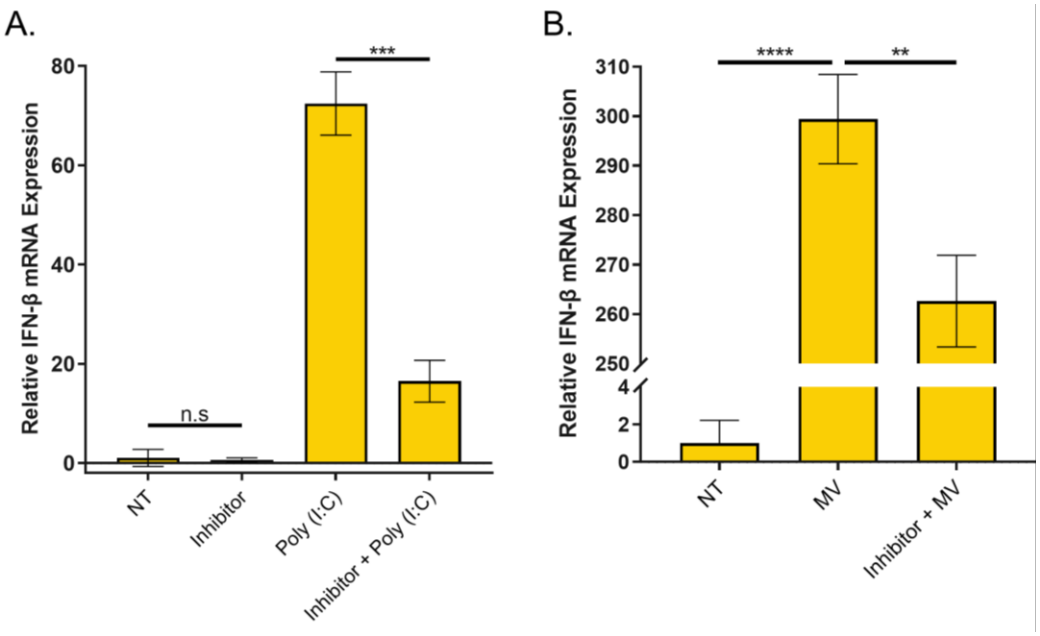


**Figure S12: RAW 264.7 cells treated with TLR3/dsRNA inhibitor produce significantly less IFN-β mRNA in response to MVs.** (A) Macrophages were pre-treated with 0.1% DMSO (NT control), HEPES-NaCl (Poly I:C control), or 10 µM TLR3/dsRNA Complex Inhibitor for 30 min at 37˚C, 5% $CO_2$. Following pre-treatment, macrophages received HEPES-NaCl (TLR3 Inhibitor control) or 10 µg/mL Poly (I:C). (B) Macrophages were pre-treated with 0.1% DMSO (NT), HEPES-NaCl (MV), or 10 µM TLR3/dsRNA Inhibitor for 30 min at 37˚C, 5% $CO_2$. Following pre-treatment, macrophages were incubated with MVs (5 µg/mL). After 3 h, total RNA was extracted and processed for β-actin and IFN-β mRNA quantitation. The data is represented as mean ±SE, n = 3. Statistical analysis was performed by one-way ANOVA with Tukey's multiple comparison test (**$p < 0.005$; ***$p < 0.001$; ****$p < 0.0001$).

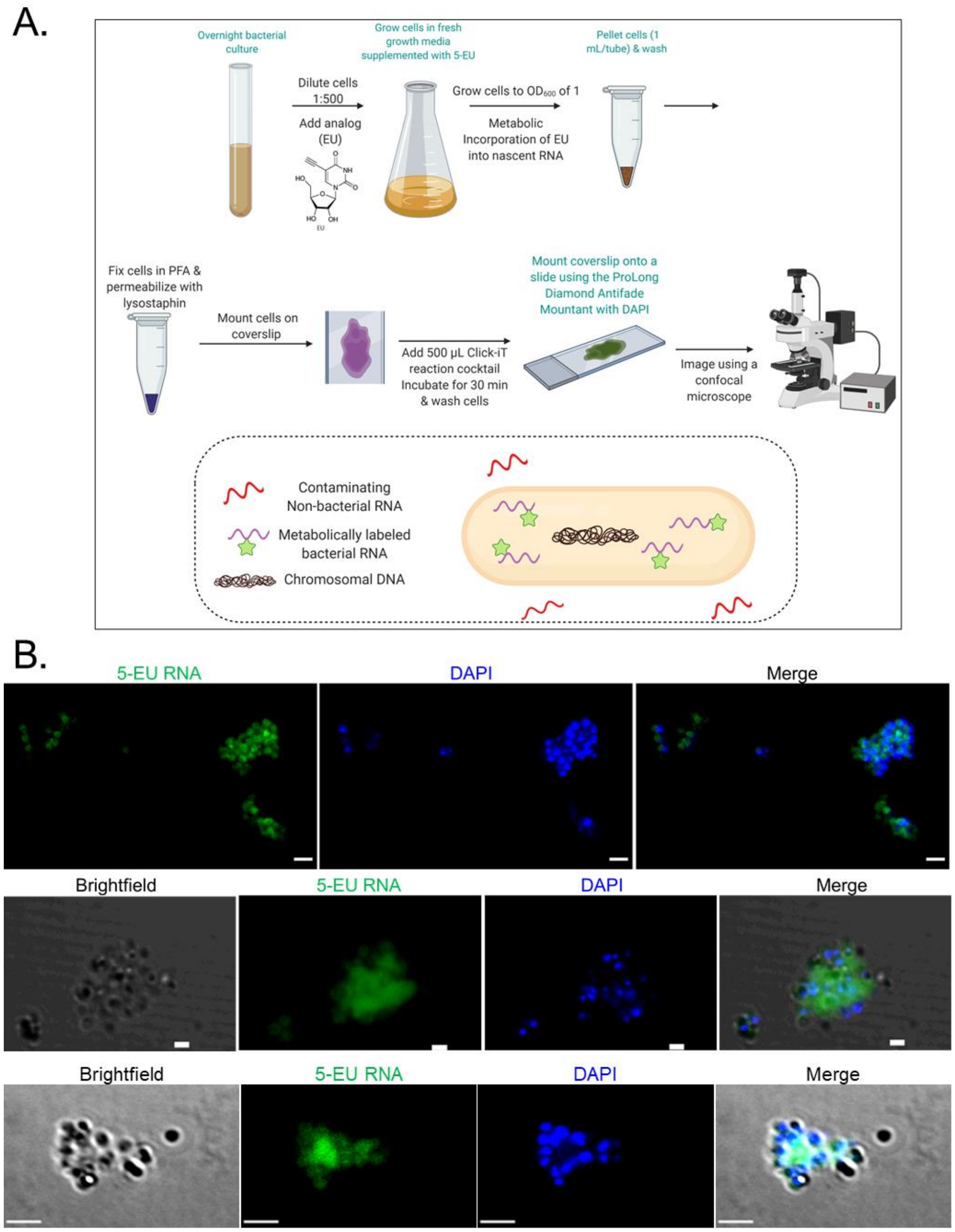


**Figure S13: 5-EU incorporation into RNA in *S. aureus* cells.** (A) Schematic representation of the metabolic labeling process for bacterial RNA. 5-EU provided in the growth media becomes incorporated into nascent RNA in actively dividing cells. At mid-log phase, cells are rinsed, fixed, and permeabilized. Fixed cells are then incubated with a fluorescent azide in the presence of a copper catalyst to enable detection of the fluorophore-azide bound to the metabolically labeled RNA using a confocal microscope. Created with BioRender.com. (B) Cells were grown in the presence of 100 µM 5-EU to $OD_{600}$ of 1. The cells were fixed, permeabilized with lysostaphin (80 µg/mL), and reacted with Alexa Fluor 488-Azide. Cells were counterstained using DAPI. Two representative images from 2 independent experiments are shown. Scale bars = 2 µm (upper and bottom panels) and 1 µm (middle panels).

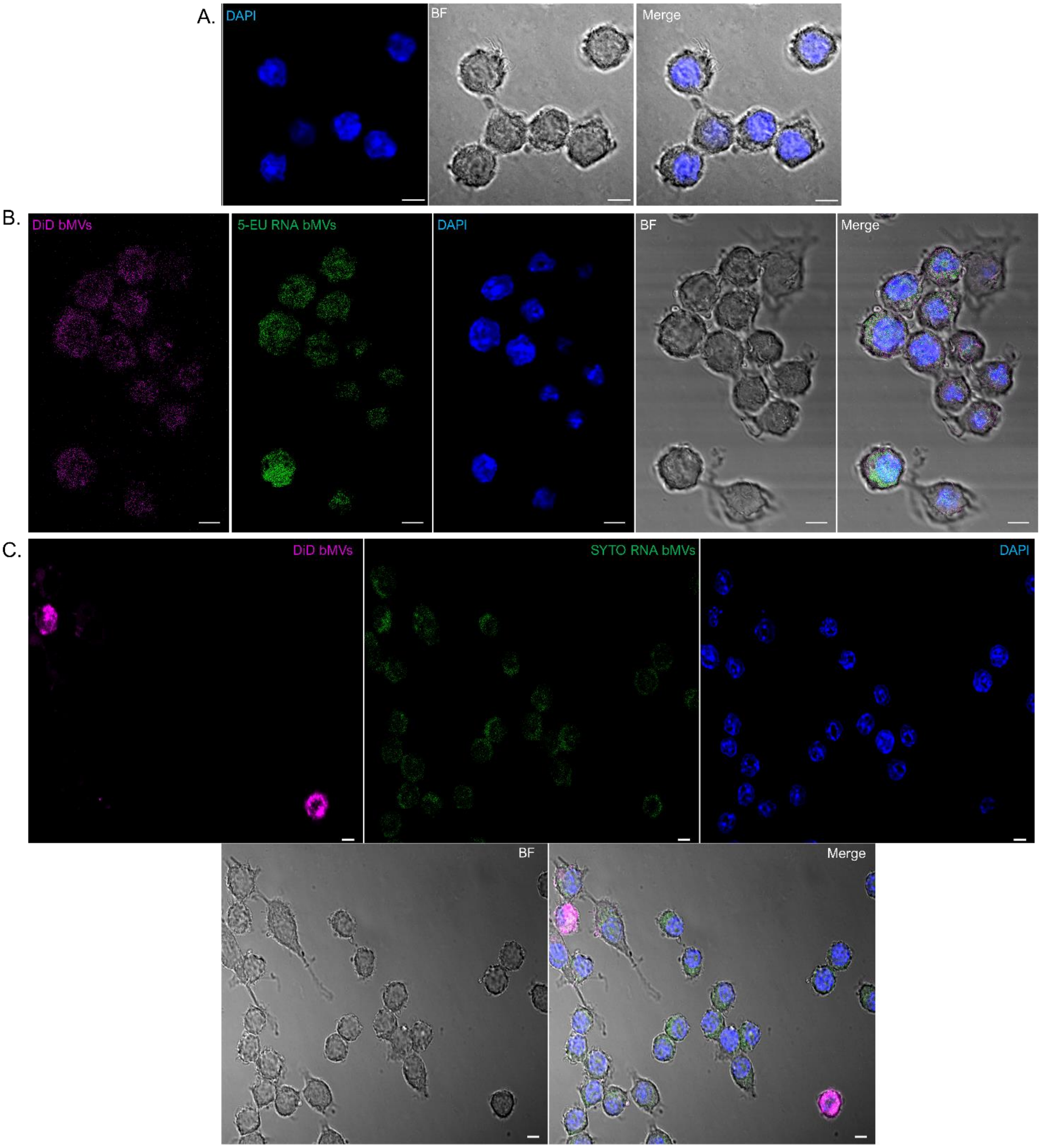


**Figure S14: Confocal microscopy images showing uptake of bMVs and bMV-RNA in wild-type macrophages.** Wild-type macrophages were treated with (A) unlabeled bMVs (5 µg/mL), (B) 5-EU RNA -DiD-bMVs (5 µg/mL), or (C) SYTO RNA-DiD-bMVs (5 µg/mL) for 1 h, fixed, and permeabilized. Macrophages treated with unlabeled bMVs or with SYTO RNA-DiD bMVs were then stained with DAPI. Macrophages treated with 5-EU RNA DiD-bMVs were reacted with the Click-iT RNA labeling kit after cell permeabilization, followed by staining with DAPI. Images are representative of 2 independent experiments. Scale bars = 5 µm.

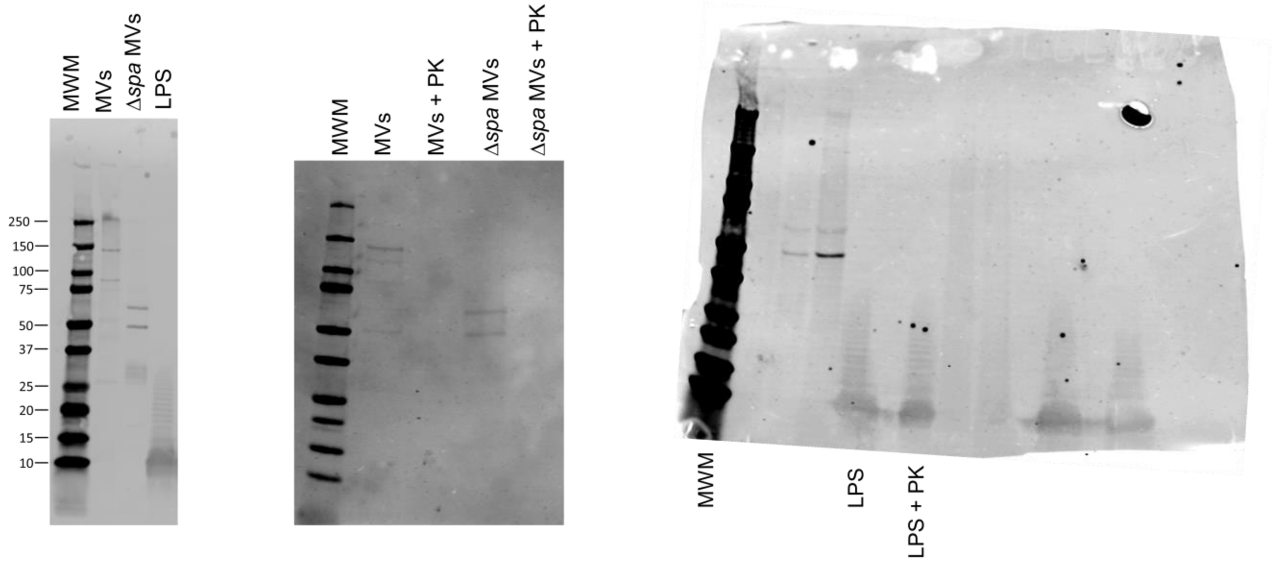


**Figure S15: Full-length western blots of images presented in Figure S5.** The labeled lanes were cropped for clarity. MWM, molecular weight marker.

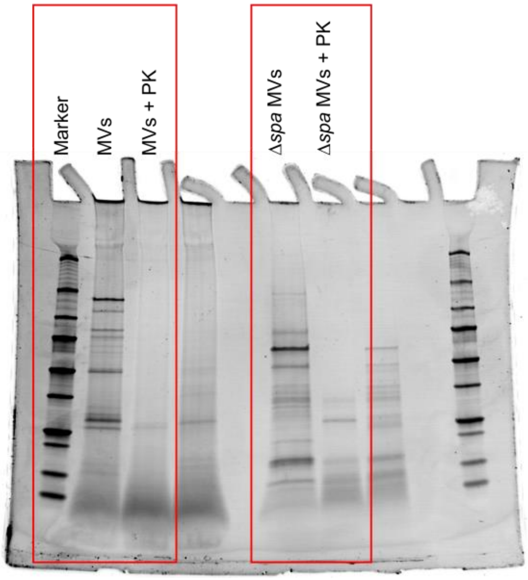


**Figure S16: Full-length SDS-PAGE gel of the image presented in Figure S6.** The labeled lanes were cropped for clarity.

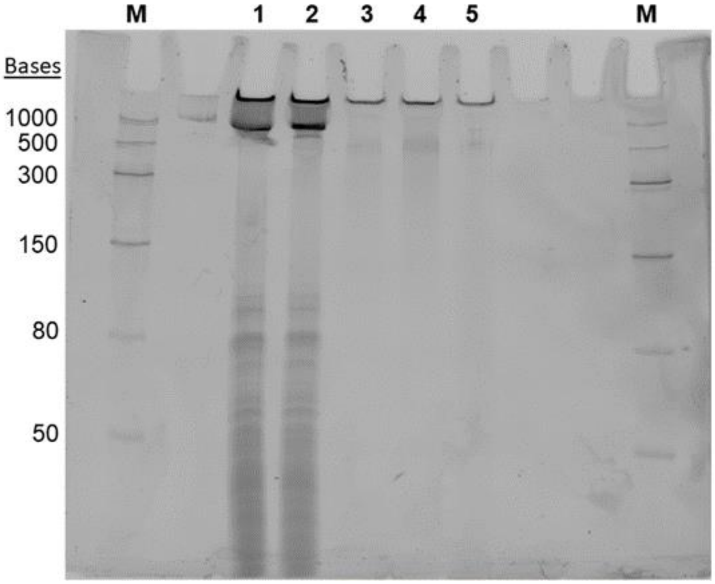


**Figure S17: Full-length image of the urea-PAGE gel presented in Figure 3.** The lanes labeled 1-5 and the marker on the left were cropped for clarity. M = Low range ssRNA ladder.

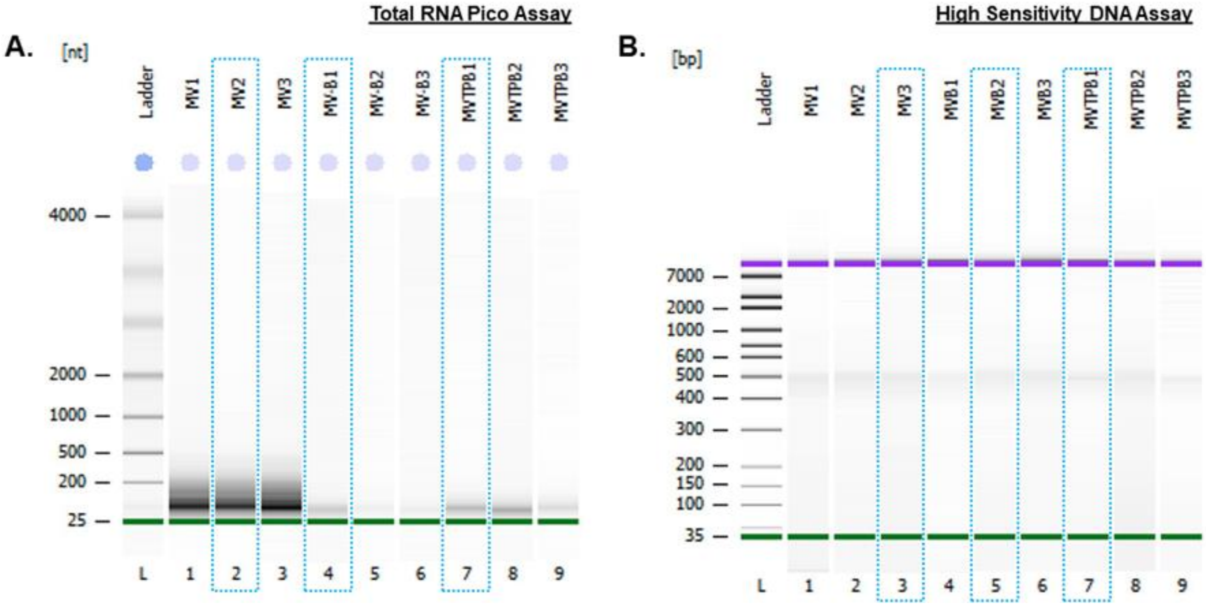


**Figure S18: Complete bioanalyzer gel images of the cropped gels presented in Figure 4B and 5B.** (A) Bioanalyzer gel image of RNA extracted from MV samples treated under different conditions. RNA from each condition was run in triplicate. Blue dotted rectangles indicate the lanes that were selected as representative images for Figure 4B. MV1 - MV3 = RNA from untreated MVs; MV-B1 – MV-B3 = RNA from bMVs; MVTPB1 - MVTPB3 = RNA from bMVs treated with Triton X-100 and Proteinase K. (B) Bioanalyzer gel image of DNA extracted from MV samples treated under different conditions. DNA from each condition was run in triplicate. Blue dotted rectangles indicate the lanes that were selected as representative images for Figure 5B. MV1 - MV3 = DNA from untreated MVs; MVB1 – MVB3 = DNA from bMVs; MVTPB1 - MVTPB3 = DNA from bMVs treated with Triton X-100 and Proteinase K.
